# Integrating genome-scale metabolic models into multi-scale simulations

**DOI:** 10.64898/2026.09.02.748294

**Authors:** Marco Ruscone, Othmane Hayoun-Mya, Randy Heiland, Paul Macklin, Alfonso Valencia, Miguel Ponce-de-Leon

## Abstract

Genome-scale metabolic models can predict how individual cells allocate resources and respond to their environment, yet few frameworks link single-cell metabolism to the spatial organisation of multicellular systems. Here we introduce PhysiCelldFBA, an extension of the PhysiCell agent-based framework that couples genome-scale dynamic flux balance analysis to off-lattice multicellular simulations. Each simulated cell carries its own metabolic model, allowing local environmental conditions to shape metabolism while metabolic activity feeds back on the surrounding environment, cellular behaviour, and spatial organisation. We first validate this coupling by showing that glucose consumption, *CO*_2_ production, and biomass accumulation remain mass-balanced in a closed E. coli system, with simulated biomass agreeing with analytical predictions to within 1%. We then demonstrate how metabolic phenotypes emerge from this coupling across microbial and mammalian systems. Spatial nutrient gradients generate metabolic stratification and acetate cross-feeding in growing E. coli colonies; diffusion-limited metabolism produces proliferative, hypoxic, and necrotic zones across a broad panel of metabolites in a tumour-like tissue; distinct, organism-specific metabolic networks give rise to syntrophic cross-feeding and spatial niche formation in a two-species consortium; and metabolic state couples energy availability to transitions between cellular motility and growth. Across these examples, metabolic stratification, cross-feeding, and phenotypic adaptation emerge from local metabolic optimisation and environmental feedback rather than being explicitly prescribed. PhysiCelldFBA therefore provides a general framework for simulating genome-scale metabolism at single-cell resolution and linking intracellular metabolic state to cellular behaviour and emergent organisation across scales.

## Introduction

Understanding how intracellular metabolic states shape population-level behaviour is a central challenge in systems biology, with implications ranging from microbial ecology to tissue homeostasis and cancer progression [1]. Genome-scale metabolic models (GSMs), together with constraint-based modelling approaches such as flux balance analysis (FBA), provide a powerful framework for predicting cellular metabolic phenotypes under physicochemical constraints [2, 3]. Advances in genome reconstruction and the integration of transcriptomic and proteomic data have enabled the generation of increasingly accurate, context-specific metabolic models for diverse organisms and tissues [4, 5]. More recently, single-cell and spatial transcriptomic technologies have opened new opportunities to investigate metabolic heterogeneity at cellular resolution [6, 7]. Together, these developments have established genome-scale metabolic modelling as a key approach for studying cellular metabolism across a wide range of biological systems.

Despite these advances, most GSM analyses rest on two assumptions that limit their applicability to spatially structured systems. The first is metabolic steady state under fixed environmental conditions: standard FBA solves a single optimisation problem for a given set of constraints, producing a flux distribution that does not evolve as the extracellular environment changes. Dynamic FBA (dFBA) relaxes this limitation by solving the FBA problem repeatedly over discrete time intervals, updating substrate availability constraints at each step from the current extracellular concentrations [8]. This allows fluxes and growth rates to evolve as conditions change—for instance, enabling a cell to switch from one carbon source to another metabolism as the first becomes depleted. The second assumption is a well-mixed environment, in which cells experience identical extracellular conditions. This precludes the nutrient gradients, cell–cell interactions, and environmental feedback that generate spatial heterogeneity in real biological systems. Spatially resolved dFBA frameworks address this limitation by coupling dynamic metabolic optimisation to an explicit representation of the extracellular environment. Platforms such as COMETS [9, 10] have demonstrated how spatial diffusion and genome-scale metabolism can predict microbial community assembly, cross-feeding and colony organisation, while agent-based approaches such as BacArena [11] represent individual cells carrying their own metabolic models on a discretised lattice. More recent developments have extended these concepts to porous environments, spatial transcriptomic data and accelerated surrogate models [7, 12–16]. Collectively, these studies have demonstrated the importance of spatially resolved metabolism in both microbial and multicellular systems.

However, most existing frameworks remain built around discretised spatial representations in which both cells and their environment are constrained by an underlying computational lattice. Although highly successful for many applications, lattice-based approaches can limit the representation of continuous cell mechanics, growth, division and spatial rearrangement, particularly in dense microbial colonies, developing tissues and tumour microenvironments. Off-lattice agent-based models offer a complementary framework by representing cells as discrete physical objects that interact mechanically while exchanging metabolites through a continuously resolved reaction-diffusion environment [17, 18]. This separation between cellular mechanics and environmental discretisation enables realistic multicellular dynamics without constraining cell positions to predefined lattice locations, making off-lattice models particularly well suited for studying the interplay between metabolism, mechanics and tissue organisation.

PhysiCell is an open-source, off-lattice agent-based framework that combines mechanically interacting cells with an efficient reaction-diffusion solver capable of modelling multiple extracellular substrates in three-dimensional tissues [19, 20]. Its modular architecture has supported a growing ecosystem of extensions incorporating intracellular signalling, extracellular matrix mechanics, pharmacokinetic/pharmacodynamic models and executable cell behaviour rules [21–25], while its computational performance has enabled applications ranging from tumour biology to viral infection and multiscale therapeutic modelling [26, 27]. Despite this mature ecosystem, PhysiCell currently lacks native support for genome-scale metabolic modelling, preventing intra-cellular metabolism from evolving dynamically alongside cell mechanics and the extracellular microenvironment.

Here we present *PhysiCelldFBA*, an open-source extension that integrates dynamic flux balance analysis into the PhysiCell framework, enabling genome-scale intracellular metabolism, continuous cell mechanics, and extracellular transport to evolve simultaneously within off-lattice, three-dimensional multicellular simulations. Rather than developing a specialised simulator for a single biological application, PhysiCelldFBA provides a general modelling framework that integrates with the existing PhysiCell ecosystem and can be applied across diverse biological systems. We first validate the numerical consistency of the metabolism-mechanics coupling through analytical mass-balance calculations, confirming quantitative agreement between extracellular substrate consumption, intracellular metabolic fluxes, and biomass accumulation. We then demonstrate the versatility of the framework across five representative applications: metabolic switching in *E. coli*, spatial metabolic organisation and cross-feeding in an expanding bacterial colony, metabolic stratification in a tumour microenvironment, cross-feeding within a syntrophic microbial consortium, and metabolism-driven cell motility. Together, these examples show that PhysiCelldFBA provides a unified platform for investigating how intracellular metabolism interacts with the physical and chemical microenvironment across scales, from single bacterial cells to multicellular tissues.

## Results

In this section, we present the results obtained with PhysiCelldFBA, including both validation of the framework and a series of use cases that demonstrate its capabilities and potential applications. We begin with a closed-batch simulation of a single bacterium to verify that model predictions are quantitatively consistent with mass conservation as well as analytical estimates of growth yield. We then describe five applications of increasing complexity, chosen to span a broad range of biological scales and systems. From single-species microbial physiology and multispecies microbial ecology to mammalian tumour biology, each addresses a biologically relevant question while collectively illustrating the versatility of the framework.

### Validation of mass conservation and flux consistency

To validate the coupling between microenvironment, metabolic uptake and secretions, and agent-based growth, we simulated the expansion of a single *E. coli* cell in a closed glucose-limited environment (see Materials and Methods). The theoretical biomass supported by the available glucose was calculated from the biomass yield predicted by the underlying genome-scale metabolic model using FBA (see Supplementary Methods S4) and compared with the biomass accumulated during the PhysiCelldFBA simulation.

The results showed that glucose decreased monotonically and was completely exhausted after approximately 3.5 h (Figure 1A). Over the same period, approximately one-third of the carbon supplied as glucose was released as CO_2_, consistent with the remainder being incorporated into biomass. As substrate was consumed, total biomass increased continuously until glucose depletion, punctuated by discrete cell-division events as the population expanded, reaching 4.09 pg dry weight when non-growth-associated ATP maintenance was not considered (Figure 1B). The analytical calculation predicted a total biomass of 4.08 pg (see Supplementary Methods S4), in strong agreement with the simulated value of 4.09 pg (relative error <1%). Similarly, when non-growth-associated ATP maintenance was included, the analytical prediction (3.87 pg) closely matched the simulated biomass (3.88 pg), again with a relative error below 1%. Equivalent agreement was obtained using the more comprehensive *E. coli* genome-scale metabolic model iML1515, demonstrating that the coupling remains robust across different metabolic reconstructions and energy maintenance constraints.

**Figure 1:**
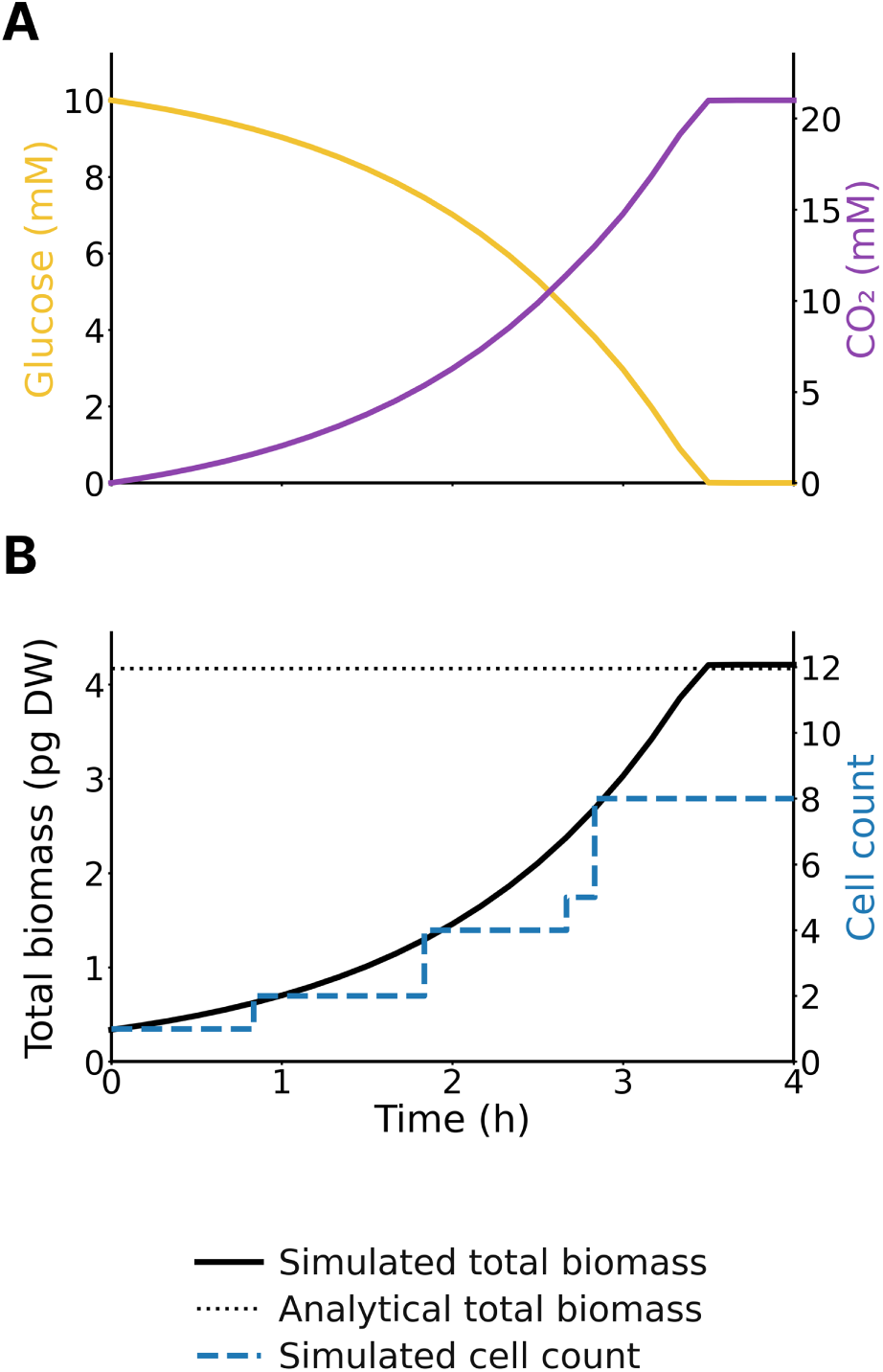
Mass conservation validation. (A) Glucose depletion and CO_2_ accumulation over time in a closed batch simulation. (B) Total biomass accumulated during the simulation (solid line, left axis) compared with the analytical prediction (dotted horizontal line), calculated from the initial glucose available in the domain and the biomass yield predicted by the underlying metabolic model; the right axis shows the corresponding cell count over time, reflecting the discrete division events.

Together, these results validate the dry-weight scaling procedure and confirm that substrate uptake, intracellular flux balance calculations, biomass accumulation, and microenvironment concentrations remain quantitatively consistent while conserving mass throughout the PhysiCelldFBA workflow. Having established the correctness of the metabolic-growth coupling, we next investigated whether the framework could reproduce emergent metabolic behaviours in the absence of explicitly encoded regulatory mechanisms.

### Emergent metabolic adaptation: the *E. coli* acetate switch

As a first use case, we simulated the acetate switch, a well-characterised metabolic transition in which *E. coli* shifts strategy in response to changing oxygen and carbon availability. This scenario tests the framework’s ability to reproduce emergent metabolic switches without explicitly encoding regulatory mechanisms, relying instead on repeated dFBA optimisation under evolving extracellular conditions. In the underlying metabolic model, acetate uptake remained dynamically coupled to local extracellular acetate concentrations through the PhysiCelldFBA transport model (see Materials and Methods).

The simulation reproduces several sequential metabolic regimes driven by the coupled depletion and resupply of oxygen (Figure 2). During an initial aerobic phase (0-1.2 h), cells grew on glucose with negligible acetate production while both oxygen and glucose remained available. As oxygen became limiting around *t*_1_ (*t ≈* 1.2 h), fermentative metabolism began, and acetate accumulated steadily as glucose consumption continued, until glucose was nearly exhausted at *t*_2_ (*t ≈* 3.4 h). Oxygen was resupplied only minutes later, at *t*_3_ (*t* = 3.5 h), leaving no sustained period of metabolic quiescence: cells switched almost immediately to aerobic acetate consumption, resuming growth and CO_2_ production until *t*_4_ (*t ≈* 5.5 h), when acetate was substantially depleted and oxygen again became limiting; the system then reached a final plateau for the remainder of the plotted window (5.5-6 h). Notably, none of these transitions was prescribed through regulatory rules; each metabolic switch, including the shift into fermentation at *t*_1_ and the near-immediate re-activation of acetate uptake at *t*_3_, emerged directly from repeated metabolic optimisation under changing environmental constraints.

**Figure 2:**
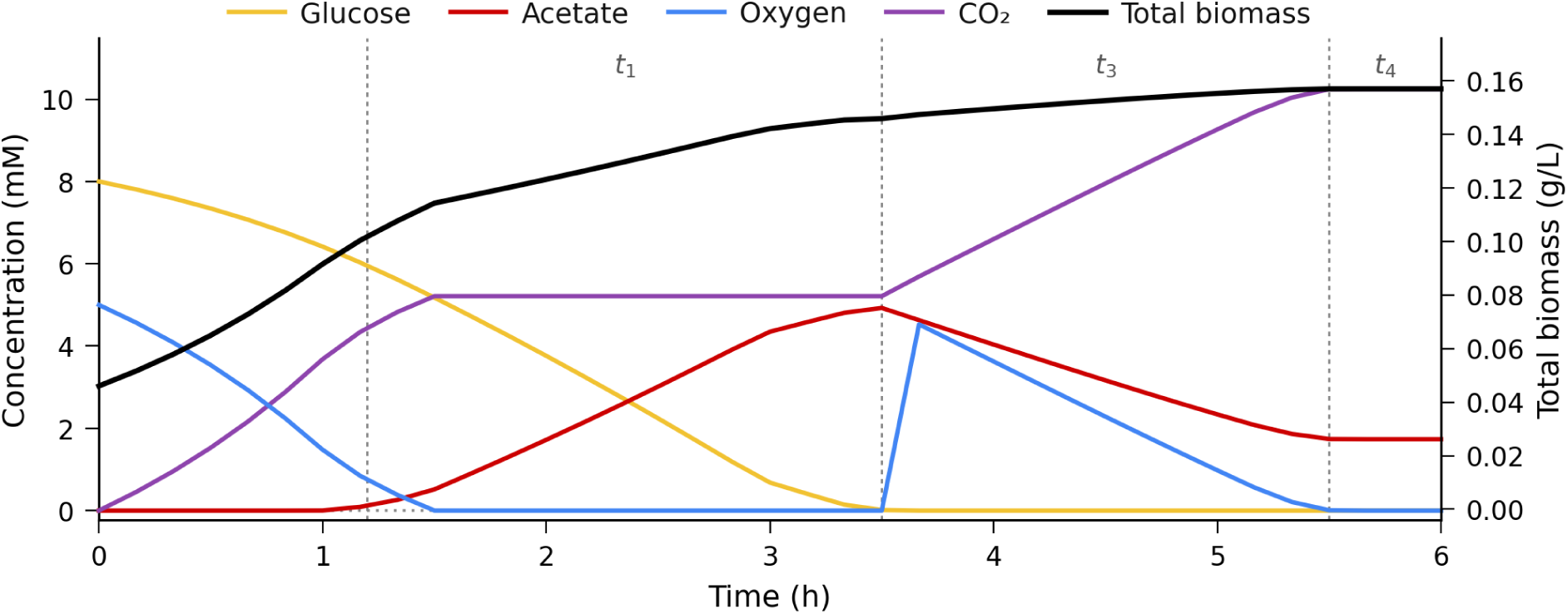
Acetate switch dynamics in *E. coli*. Coloured lines indicate the temporal evolution of extracellular glucose, oxygen, acetate, CO_2_, and total biomass over the course of the simulation. Dashed vertical lines mark the transitions between metabolic phases, at *t*_1_ *≈* 1.2 h, *t*_3_ = 3.5 h, and *t*_4_ *≈* 5.5 h (*t*_2_ *≈* 3.4 h, glucose exhaustion, is not marked separately as it coincides almost exactly with *t*_3_). An initial aerobic phase (0–*t*_1_) gives way to oxygen-limited fermentation and acetate secretion (*t*_1_–*t*_2_); glucose exhaustion at *t*_2_ is followed almost immediately by an oxygen resupply at *t*_3_, leaving only a brief transition before cells switch to aerobic acetate re-utilisation and renewed growth (*t*_3_–*t*_4_), reaching a final plateau (*t*_4_–6 h) as oxygen again becomes limiting.

This temporal switch was accompanied by a corresponding spatial reorganisation of extracellular metabolites (Supplementary Figure S5): glucose gradients were shallow during the initial aerobic phase (0-1.2 h), acetate accumulation produced outward, source-like gradients during the fermentative phase (1.2-3.4 h), and by the oxygen-resupply phase (3.5-5.5 h) acetate had become the dominant carbon source, reversing its earlier role from secreted by-product to primary substrate. Together, these temporal and spatial dynamics demonstrate that coupling metabolic optimisation with cellular transport and a dynamically evolving environment is sufficient to generate adaptive metabolic transitions without predefined regulatory mechanisms.

### Spatial metabolic organisation in a growing *E. coli* colony

We next simulated an expanding *E. coli* colony cultured in minimal medium with glucose as the sole carbon source under aerobic conditions, to test whether the framework could resolve metabolic heterogeneity emerging purely from spatial structure within an initially homogeneous, clonal population. We seeded an initial colony and simulated an *in silico* growth experiment for 10 hours (see Materials and Methods). The colony underwent an initial phase of near-exponential growth during the first 5 hours, after which total biomass accumulation progressively decelerated as the colony interior became nutrient-limited (Figure 3A); this transition from exponential to sub-exponential growth kinetics is shown more clearly on a logarithmic scale in Supplementary Figure S6.

**Figure 3:**
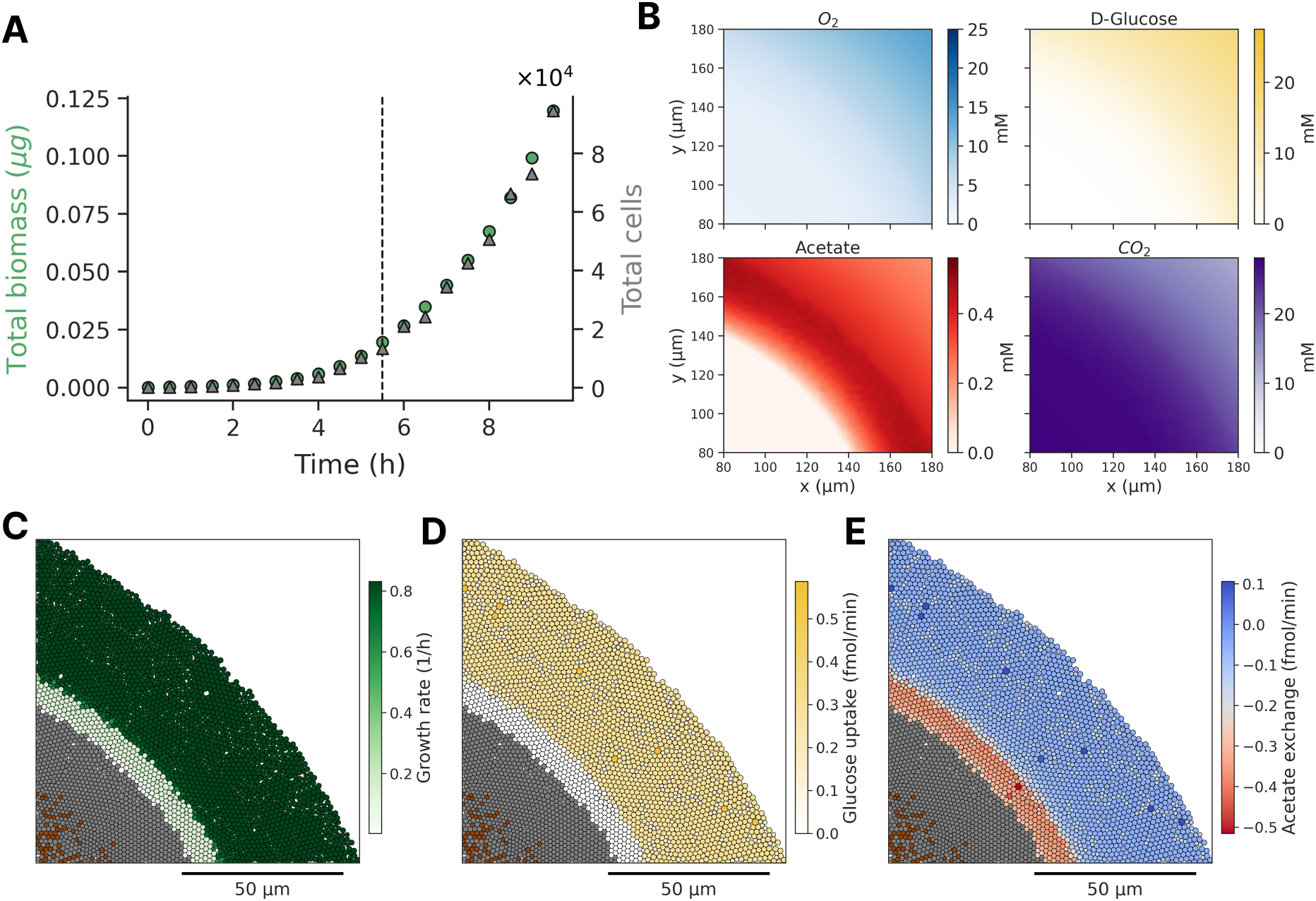
Growth dynamics and spatial metabolic organisation in an expanding *E. coli* colony. (A) Temporal evolution of total biomass showing an initial exponential growth phase during the first 5 h. (B) Spatial distribution of extracellular metabolites at 9 h, including oxygen (*O*_2_), D-glucose, acetate, and *CO*_2_. (C–E) Single-cell metabolic states mapped onto the spatial domain. Grey cells represent dead cells. (C) Growth rate distribution (1/h), showing a peripheral region of fast-growing cells, an intermediate region of reduced growth, and a central region of non-viable cells. (D) Glucose uptake rates highlight limited substrate availability in the colony interior. (E) Acetate exchange fluxes indicate spatial variation in secretion and uptake across the population.

By 9 hours, oxygen and glucose were substantially depleted in the colony interior while remaining available at the periphery (Figure 3B), and this extracellular gradient was directly reflected in cellular phenotype: cells at the periphery maintained high growth rates and active glucose uptake, cells in intermediate regions showed progressively reduced proliferation, and a central core of non-viable cells emerged from sustained nutrient limitation (Figure 3C–D), consistent with the formation of inactive cores reported in dense microbial colonies [28]. Acetate exchange fluxes revealed a further layer of spatial structure: peripheral cells secreted acetate as a by-product of overflow metabolism, while inner cells showed reduced secretion and net acetate re-assimilation (Figure 3E and Supplementary Video S1), establishing a spatial division of metabolic labour generated solely by local differences in resource availability, with no phenotype imposed *a priori*. The spatially segregated acetate cross-feeding pattern and its relationship to comparable findings in the literature are further discussed in the Discussion section. We next investigate the emergence of spatial metabolic stratification in a simulated population of cancer cells growing near a nutrient source.

### Metabolic stratification in a tumour-like microenvironment

We implemented a model in which MCF7 cells were arranged adjacent to a blood vessel serving as the source of oxygen and nutrients, with cellular metabolism represented by a genome-scale reconstruction tailored to the cell line and constrained by exometabolomic data [29], enabling the simulation of uptake and secretion fluxes across a broad panel of extracellular metabolites (see Materials and Methods).

Spatial simulation recovered a clear gradient in cellular growth rate with respect to blood vessel distance, giving rise to three metabolically distinct zones: a proliferative region (0-120 *µ*m) with active biomass production, a hypoxic intermediate zone (120-200 *µ*m) of progressively reduced growth, and a necrotic core beyond 200 *µ*m in which intracellular fluxes fell below maintenance requirements (Figure 4A). Critically, this zonation was recovered simultaneously across the full panel of metabolites included in the reconstruction, not only oxygen and glucose: Figure 4B (with the corresponding extracellular concentration profiles shown in Supplementary Figure S8) shows coherent, zone-consistent uptake and secretion profiles for fifteen metabolites, including multiple amino acids (glutamine, glycine, threonine, isoleucine, leucine, lysine, serine, phenylalanine), nitrogen and phosphate handling (ammonium, phosphate, ornithine), and core fermentative by-products (CO_2_, lactate), all varying systematically with distance from the vessel as a direct consequence of each metabolite’s independent diffusion and consumption dynamics rather than any explicit zonation rule [30]. The resulting boundaries are consistent with the broad range of diffusion-limited zonation reported for solid tumours [31–33].

**Figure 4:**
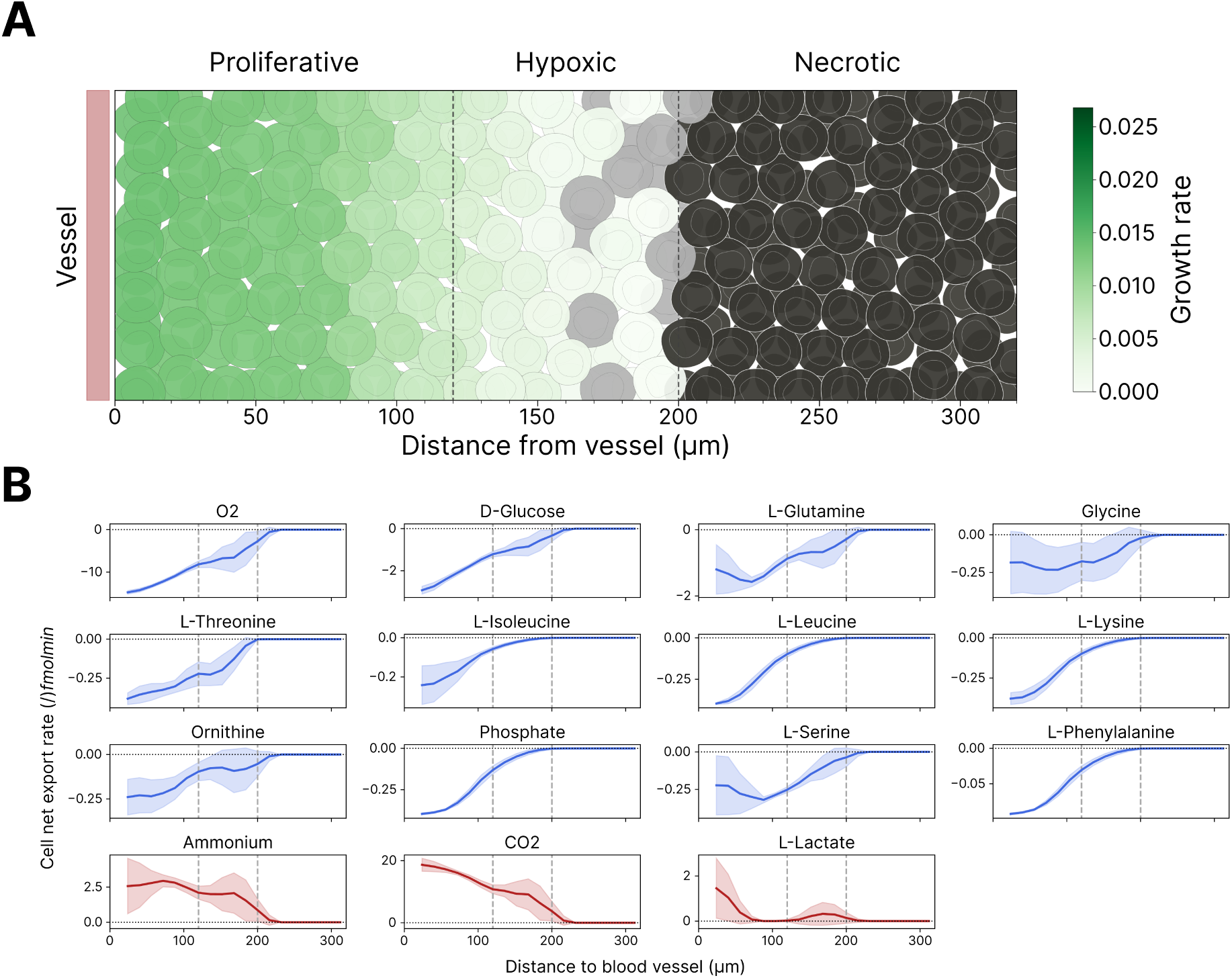
Growth rate and viability distribution in a tumour-like cell population near a blood vessel, indicated as a red rectangle. **(A)** Spatial distribution of cells arranged adjacent to a blood vessel (left), which supplies nutrients and oxygen. Colours indicate cell state (alive, pre-dead, dead), while the green intensity represents growth rate (1/h). The dotted lines indicate proliferative, hypoxic, and necrotic regions defined by oxygen and glucose availability. **(B)** Profiles of single-cell net uptake and secretion rates for multiple metabolites as a function of distance to the vessel; negative and positive values correspond to uptakes and secretions, respectively, and shaded regions indicate variability across cells.

Beyond the population-averaged trends shown in Figure 4B, single-cell maps of metabolite uptake and secretion (Supplementary Figure S9, Supplementary Videos S2 and S3) reveal substantial cell-to-cell heterogeneity in oxygen, glucose, lactate, and glutamine flux even within the proliferative zone, with a subset of cells consistently showing markedly higher flux than their immediate neighbours despite sharing the same local environment.

Together, these results show that PhysiCelldFBA can efficiently simulate physiologically realistic microenvironments comprising many diffusing metabolites, reproducing the spatial metabolic stratification that arises during tumour growth. We next asked whether the framework could similarly capture metabolic interactions between distinct organisms, each governed by its own genome-scale network.

### Emergent cross-feeding in a synthetic microbial consortium

To illustrate how the framework extends to multispecies communities, we implemented a syntrophic consortium composed of the fermentative bacterium *Clostridium beijerinckii* (CB) and the methanogenic archaea *Methanosarcina barkeri* (MB), coupled through the exchange of hydrogen and acetate (Figure 5A): these metabolites are released as fermentative by-products by CB and serve as the primary carbon and energy sources sustaining MB growth.

**Figure 5:**
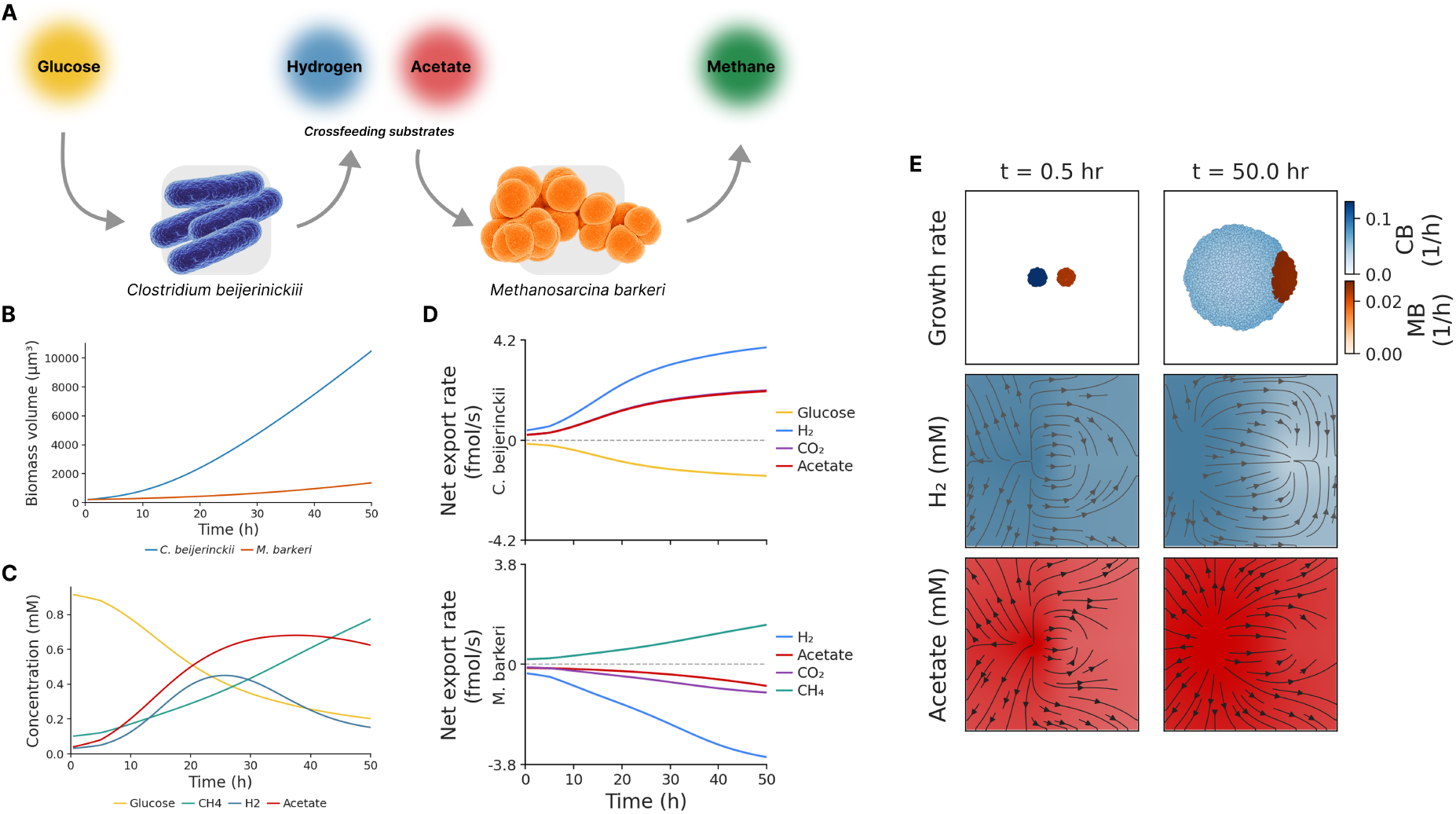
Cross-feeding dFBA model implementation and emergent dynamics. (A) Schematic of the syntrophic system composed of C. beijerinckii (CB; left) and M. barkeri (MB; right). CB consumes glucose and releases hydrogen and acetate as fermentative byproducts; MB can consume hydrogen and CO_2_ via hydrogenotrophic methanogenesis or only acetate via acetoclastic methanogenesis, producing methane. (B) Biomass volume (*µ*m^3^) trajectories for both populations over the course of the simulation, showing quasi-exponential CB growth under continuous glucose supply and a slower but sustained MB expansion driven by CB-derived metabolites. (C) Total population net export rate (fmol/s) for CB (top) and MB (bottom). The top panel shows CB trends: net glucose uptake (negative, yellow) alongside H_2_ and acetate export (positive, blue and red); the bottom panel shows MB trends: net H_2_ and CO_2_ uptake growing in magnitude alongside progressive CH_4_ export (green). (D) Population-level total substrate exchange (mmol per dFBA time step) summed across all cells of both species. The top panel shows depletion of glucose; the second panel shows continuous methane production, and the bottom panels show the diffusion dynamics of cross-feeding substrates, CB-driven accumulation and then consumption by the MB population. (E) Spatial microenvironment and cell population snapshots at 0.5 h and 50 h simulation time points with arrows indicating the concentration gradients.

We simulated the consortium for 50 h (see Materials and Methods). Both populations demonstrated sustained growth throughout the simulation but with markedly different kinetics (Figure 5B). CB expanded rapidly under direct access to the supplied glucose, while MB grew more slowly, consistent with the lower growth yield of methanogenesis and its dependence on CB-derived substrates. The full spatiotemporal evolution of growth rate across the simulated domain is shown in Supplementary Video S4, with sequential snapshots shown in Supplementary Figure S10. As expected from this metabolic coupling, CB progressively released H_2_ and acetate, which MB consumed alongside CO_2_ to produce methane (Figure 5C–D); Hydrogen uptake dominated the methanogen flux balance overall, but acetate consumption increased over the course of the simulation, indicating a mixture of hydrogenotrophic and acetoclastic activity rather than reliance on a single pathway.

Comparison of the spatial substrate fields at the start and end of the simulation (Figure 5E) showed that substrate gradients progressively emerged as the community grew. By the end of the simulation (50 h), CB dominated the biomass and generated a broad acetate-rich zone. At the same time, MB remained confined to a locally H_2_-depleted region at the colony periphery, with gradient directions consistent with net consumption and net production maintained by diffusion-limited exchange between the two populations. Sequential snapshots of the corresponding substrate-field dynamics across the full simulation are provided in Supplementary Figure S11. These results establish that PhysiCelldFBA can capture emergent ecological structure. Functional specialisation, resource partitioning, and spatial niche formation arise from the coupled metabolism of distinct organisms, without these interactions being explicitly parameterised. In the final use case, we extend the framework beyond growth and metabolite exchange to couple intracellular state to cell behaviour.

### Coupling intracellular metabolism to cell motility

As a final use case, we asked whether the framework could couple intracellular metabolic state to a cellular behaviour beyond growth, since many cellular behaviours impose metabolic demands that compete directly with biosynthesis. We developed a minimal model of metabolism-driven chemotaxis combining PhysiCell’s built-in motility and chemotaxis modules with intracellular dFBA, in which migration speed is determined by the ATP production capacity predicted by the metabolic network, and a hysteretic phenotypic switch governs the transition between motility and growth as a function of local glucose availability (see Materials and Methods).

Representative simulations revealed two distinct behavioural regimes emerging from the interaction between glucose availability, substrate uptake kinetics, and intracellular ATP production (Figure 6). Under high glucose supply and low *K_m_*, the cell migrated efficiently toward the glucose-rich region while sustaining near-maximal ATP-driven migration speed (Figure 6A, C, E). Once the cell reached the glucose-rich boundary, the metabolic objective switched from motility to biomass optimisation, triggering growth recovery and division of the cell; this was accompanied by a sharp drop in migration speed and motility-associated ATP flux, while glucose uptake was maintained to support the resulting daughter cells (Figure 6C, E). Increasing *K_m_*under the same high-glucose boundary condition produced a qualitatively different outcome: the cell generated sufficient ATP to sustain directed migration, with migration speed increasing gradually as it approached the glucose-rich region, but glucose uptake remained below the threshold required to activate the growth objective; consequently, motility persisted with no proliferation throughout the simulation (Figure 6B, D, F).

**Figure 6:**
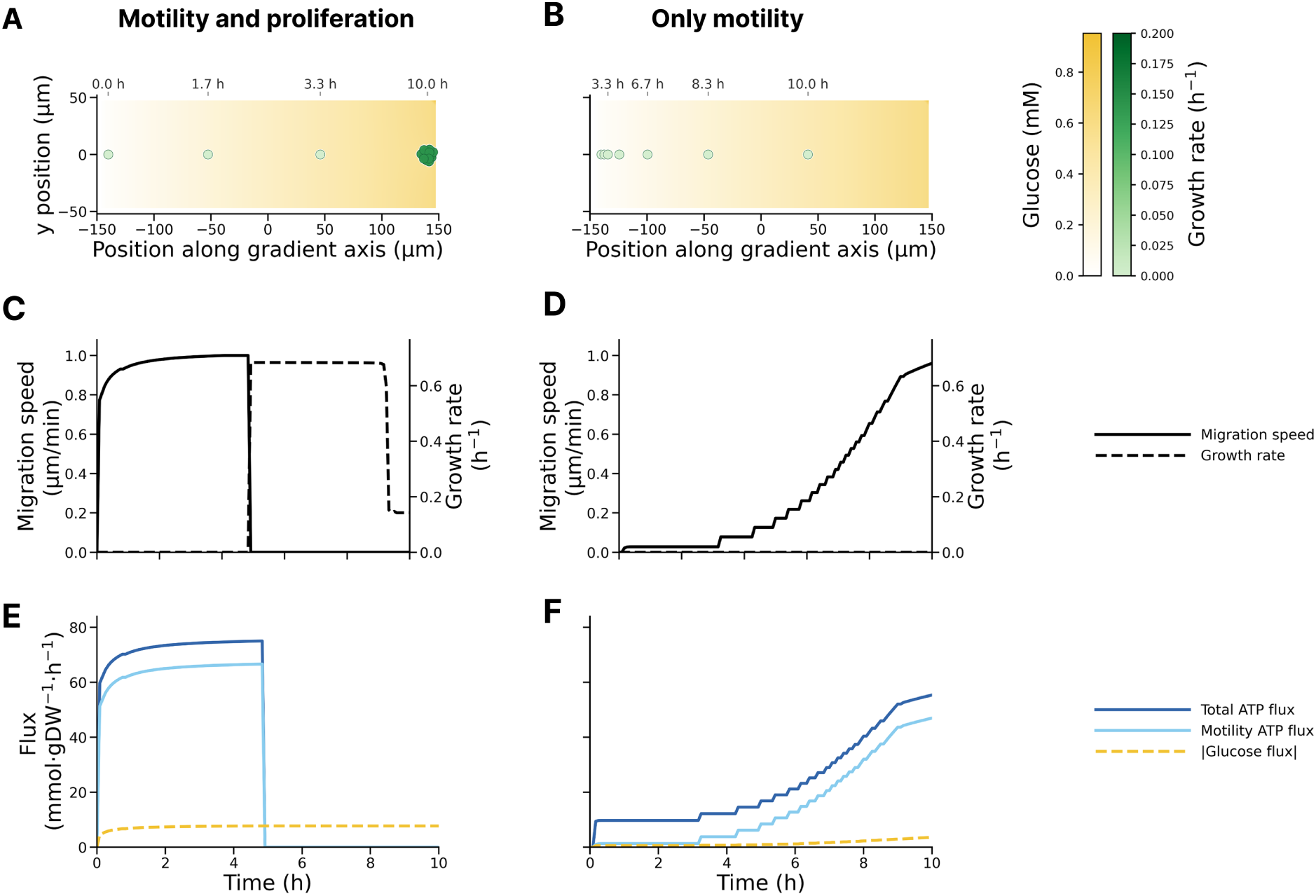
Parameter-dependent motility and proliferation under glucose limitation. (A) Spatial trajectory of a representative cell under growth-permissive conditions (low *K_m_*, Hill coefficient of 1), migrating along the glucose gradient toward the glucose-rich boundary; clustering of trajectory points near the boundary reflects the onset of cell division. (B) Spatial trajectory of a representative cell under the same glucose boundary condition but with a higher *K_m_*, showing sustained motility along the gradient without proliferation. (C-D) Migration speed (solid line) and growth rate (dashed line) over time for the growth-permissive (C) and motility-only (D) conditions. In (C), migration ceases and growth resumes once the cell reaches the glucose-rich region and divides; in (D), migration speed increases gradually as the cell approaches the boundary, while growth rate remains at zero throughout. (E-F) Corresponding metabolic flux dynamics (total ATP flux, motility-associated ATP flux, and glucose uptake flux) for the growth-permissive (E) and motility-only (F) conditions.

This use case illustrates that the framework can be extended beyond growth-focused applications to investigate the metabolic regulation of complex cellular behaviours such as motility and chemotaxis; we discuss the relationship of these phenotypic regimes to experimentally established growth-motility trade-offs in the Discussion.

## Discussion

The examples presented here demonstrate that coupling genome-scale metabolism to spatially resolved multicellular simulations provides a powerful framework for understanding how complex biological behaviours emerge from local metabolic interactions. PhysiCelldFBA extends the PhysiCell ecosystem with genome-scale metabolic modelling, enabling intracellular metabolism, extracellular transport, and multicellular mechanics to evolve simultaneously within a single spatially explicit framework. Across diverse applications spanning microbial physiology, microbial ecology, and tumour biology, our results demonstrate that coupling dynamic flux balance analysis to off-lattice agent-based modelling provides a general approach for investigating how metabolism shapes cellular behaviour across spatial and temporal scales. Rather than representing a specialised model for a particular biological system, PhysiCelldFBA provides a common computational framework in which the same underlying principles govern systems ranging from bacterial colonies to multicellular tissues.

A central observation emerging from all case studies is that complex multicellular behaviours arise naturally from the reciprocal interaction between intracellular metabolic optimisation and a dynamically changing microenvironment. Several of the behaviours reproduced here—including acetate switching, metabolic cross-feeding, and spatial metabolic zonation—were not explicitly programmed into the model. Instead, they emerged from the feedback between cellular metabolism and extracellular nutrient availability. The motility example differs in that a predefined hysteretic switch couples local glucose availability to the competing motility and growth objectives, while the metabolic state determines the energetic capacity for these behaviours.

Spatial organisation is central to this emergent behaviour. In well-mixed environments, metabolic adaptation is expressed primarily as a temporal process, as illustrated by the sequential glucose-to-acetate transition in the acetate switch model. Once space is explicitly resolved, however, the same metabolic logic gives rise to persistent spatial heterogeneity. In the microbial colony, nutrient gradients generate metabolically distinct subpopulations occupying different ecological niches, while in the tumour model, diffusion-limited substrate availability produces proliferative, hypoxic, and necrotic regions despite all cells sharing the same intracellular metabolic network. These examples illustrate that spatial heterogeneity need not arise from genetic differences or predefined cell states, but can emerge solely through local environmental constraints acting on identical metabolic networks. More broadly, they highlight PhysiCelldFBA’s ability to resolve intracellular metabolism at single-cell resolution while simultaneously capturing the physical and chemical interactions that organise multicellular systems.

The spatial division of metabolic labour predicted by the microbial colony model provides an important biological validation of the framework. The coexistence of glucose-fermenting cells at the colony periphery and acetate-consuming cells in the colony interior closely mirrors both experimental and computational observations of *E. coli* colony metabolism. In particular, ^13^C metabolic flux analysis identified a comparable partition into glucose-consuming acetate secretors and acetate-consuming subpopulations [34], while three-dimensional reaction–diffusion FBA simulations reproduced a spatially organised acetate cross-feeding architecture that was subsequently confirmed experimentally [35]. PhysiCelldFBA recovers this organisation directly at single-cell resolution as an emergent continuum of metabolic states. The close agreement between these independent experimental, computational and modelling approaches provides confidence that the coupling between intracellular metabolism, extracellular transport and cell-based dynamics captures biologically meaningful spatial organisation.

At the same time, our simulations highlight unresolved biological questions. Cole *et al.* suggested that simultaneous glucose and acetate uptake represented a modelling artefact of unconstrained FBA, motivating the incorporation of regulatory constraints to enforce an acetate switch [35]. The small co-utilising population observed in our simulations may similarly represent either a genuine transient metabolic phenotype or an artefact arising from the optimisation formulation. Likewise, recent agent-based simulations have suggested that acetate cross-feeding contributes relatively little to biomass production and instead primarily supports cellular maintenance following glucose depletion [28]. Our results are broadly consistent with this interpretation, as acetate-consuming cells exhibit substantially lower growth rates than peripheral glucose-consuming cells. These observations illustrate the framework’s ability to generate experimentally testable hypotheses while identifying aspects of microbial metabolism that remain incompletely understood.

The tumour and syntrophic consortium models demonstrate that these principles extend well beyond bacterial colony physiology. In the tumour microenvironment, physiologically realistic proliferative, hypoxic and necrotic regions emerged solely through diffusion-limited nutrient availability acting on a common intracellular metabolic network. Importantly, these transitions did not require explicit concentration thresholds or phenotype-switching rules. Instead, they arose naturally as oxygen and glucose concentrations approached the kinetic regimes in which cellular uptake became negligible. Because the underlying reconstruction is genome-scale, this spatial organisation was recovered simultaneously across a broad range of metabolites rather than being limited to a small number of manually selected nutrients. As increasingly detailed exometabolomic datasets and context-specific metabolic reconstructions become available, this capability offers a route towards spatially resolved metabolic models that more faithfully reflect the biochemical complexity of living tissues.

The syntrophic consortium extends the framework in a complementary direction by demonstrating that PhysiCelldFBA can simultaneously simulate distinct organisms carrying independent genome-scale metabolic reconstructions within a shared physical environment. Here, hydrogen- and acetate-mediated cross-feeding between *C. beijerinckii* and *M. barkeri* emerged directly from the independent optimisation of each organism’s metabolic network under locally varying environmental conditions, without prescribing a division of labour or spatial organisation. Together, the microbial colony, tumour and consortium examples demonstrate that PhysiCelldFBA is not tailored to a particular organism or biological question. Instead, it provides a common mechanistic framework in which spatial metabolic organisation, ecological interactions and community structure emerge from the same underlying coupling between intracellular metabolism, extracellular transport and multicellular dynamics.

The motility example illustrates that the scope of PhysiCelldFBA extends beyond predicting growth and metabolite exchange. By coupling ATP availability directly to cell migration, intracellular metabolic state becomes a regulator of higher-level cellular behaviour, giving rise to distinct phenotypic strategies characterised by growth, persistent motility, or metabolic quiescence. Although this model was intentionally simplified and was not designed as a quantitative description of bacterial chemotaxis, the resulting growth–motility trade-offs are qualitatively consistent with experimental observations that bacterial investment in motility depends strongly on energetic status and nutrient availability. More broadly, this example demonstrates that genome-scale metabolism can serve as a mechanistic link between environmental conditions and diverse cellular decision-making processes, complementing the behavioural modules already available within PhysiCell.

From a methodological perspective, the behaviour observed across all case studies is underpinned by the explicit coupling between intracellular metabolism and extracellular transport. PhysiCelldFBA enforces local mass conservation by ensuring that cellular uptake cannot exceed the substrate available within each voxel, while metabolic uptake and secretion continuously reshape the surrounding microenvironment. This reciprocal feedback establishes a dynamic interaction in which environmental conditions constrain intracellular metabolism, metabolism modifies the extracellular environment, and the resulting environmental changes influence subsequent cellular behaviour. Notably, this coupling alone is sufficient to generate metabolic adaptation, spatial heterogeneity, ecological interactions and behaviour switching across a broad range of biological systems, without requiring these phenomena to be encoded explicitly. The analytical mass-conservation test presented here provides a simple but rigorous validation of this coupling while also illustrating an important distinction between continuous biomass accumulation and discrete agent-based proliferation. Although biomass production follows the analytical mass balance, cell division remains a discrete event, leaving any residual biomass distributed among the existing cells rather than creating fractional agents.

Like all agent-based implementations of dynamic flux balance analysis, PhysiCelldFBA necessarily incurs a substantial computational cost because each cell requires an independent metabolic optimisation throughout the simulation. However, this computational burden is accompanied by a high degree of intrinsic parallelism. Diffusion calculations inherit the efficient reaction–diffusion solvers developed within BioFVM, including distributed-memory implementations capable of scaling across high-performance computing systems, while the metabolic optimisation performed for each individual cell is naturally embarrassingly parallel. Together, these characteristics provide a clear path towards increasingly large multicellular simulations incorporating detailed genome-scale reconstructions as high-performance computing architectures continue to evolve.

PhysiCelldFBA also inherits several important limitations from the underlying flux balance analysis formalism. First, cellular metabolism is represented as the solution to an optimisation problem, requiring the specification of an objective function. Although biomass maximisation provides a useful approximation for many experimental systems, cells frequently balance competing objectives related to maintenance, stress adaptation, migration, persistence and other physiological processes. The motility example presented here illustrates one alternative objective based on ATP production, but future applications will likely benefit from multi-objective optimisation strategies or the incorporation of regulatory information derived from transcriptomic, proteomic or signalling data. Second, as in all constraint-based models, multiple equivalent optimal flux distributions may exist. In spatial simulations, these alternative solutions can propagate to extracellular exchange fluxes and ultimately influence emergent nutrient gradients and multicellular behaviour. Integrating regulatory constraints or context-specific omics data offers a promising route for reducing this ambiguity while further improving biological realism. More generally, PhysiCelldFBA currently captures metabolic adaptation arising from environmental constraints but does not explicitly represent intracellular regulatory processes. The ability to combine genome-scale metabolism with regulatory and signalling models already available within the PhysiCell ecosystem, such as PhysiBoSS, represents a particularly promising direction for future development.

Taken together, these results position PhysiCelldFBA as a complementary addition to the growing ecosystem of multiscale modelling tools built around PhysiCell. Existing extensions have progressively incorporated intracellular signalling, extracellular matrix mechanics, pharmacological modelling and rule-based cellular behaviours. PhysiCelldFBA introduces genome-scale metabolism as a new mechanistic layer capable of interacting directly with these existing components, providing a unified framework in which metabolism, signalling, mechanics and behaviour can evolve simultaneously within spatially resolved multicellular systems. As genome-scale reconstructions, spatial transcriptomics, metabolomics and single-cell technologies continue to mature, the ability to integrate these complementary data sources into predictive mechanistic models will become increasingly important. PhysiCelldFBA provides a foundation for this next generation of multiscale digital models by allowing metabolism to participate directly in the emergence of tissue organisation, microbial ecology and multicellular function rather than treating it as an isolated intracellular process.

## Methods

### Framework overview

PhysiCell is an open-source, off-lattice agent-based simulator in which cells are represented as physically interacting agents that move, grow, divide, and die within a continuous three-dimensional domain [19], while extracellular substrates are resolved using the BioFVM reaction–diffusion solver [20]. Here, we extend this framework with genome-scale metabolic modelling through dynamic Flux Balance Analysis (dFBA).

PhysiCelldFBA couples an intracellular metabolic model to the local extracellular microenvironment of each cell. At each dFBA update, local substrate concentrations are used to constrain the corresponding exchange reactions, and the resulting FBA problem is solved independently for each cell. The predicted metabolic state determines cellular growth and metabolite exchange, which in turn modify the extracellular environment through BioFVM, creating a bidirectional feedback between intracellular metabolism, cellular behaviour, and the microenvironment (Supplementary Figure S1).

The FBA updates are synchronised with the BioFVM diffusion timestep so that exchange constraints are calculated from the current local substrate concentrations. Uptake is limited both by concentration-dependent transport kinetics and by the amount of substrate physically available within the local voxel. This prevents numerical over-consumption and maintains local mass conservation. The detailed mathematical formulation is given below, with the complete derivations and dimensional scaling provided in Supplementary Methods S1–S3.

### Mathematical formulation

At each FBA update, the metabolic state of a cell is calculated by solving the constrained linear programme

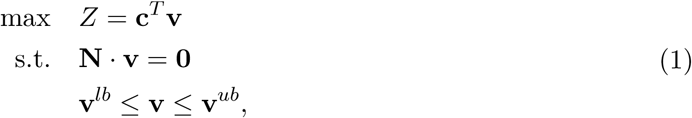

where **N** is the stoichiometric matrix of the metabolic model, **v** is the flux vector, **c** defines the objective function, and **v***^lb^* and **v***^ub^* are the reaction bounds. The objective function *Z* typically corresponds to the biomass reaction. However, alternative objectives can be specified for specialised use cases, such as ATP maximisation in the motility example presented in the Results. The extracellular microenvironment is represented by the BioFVM reaction–diffusion equation,

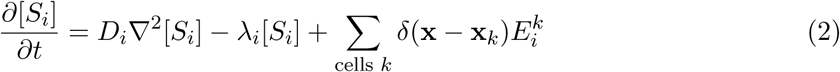

where (*D_i_*) is the diffusion coefficient, (*λ_i_*) the extracellular decay (removal) rate, and (*E^k^*) the net export rate of substrate (i) by cell (k). PhysiCelldFBA couples the two formulations by using local substrate concentrations to constrain metabolic exchange reactions and by using the resulting cell-level exchange fluxes as source or sink terms in the extracellular reaction–diffusion system.

Exchange reaction bounds are updated at each time-step from the local extracellular environment through two complementary constraints. First, uptake is limited by Michaelis–Menten kinetics based on the local substrate concentration. For each substrate *i*, the concentration-dependent kinetic uptake bound is calculated as

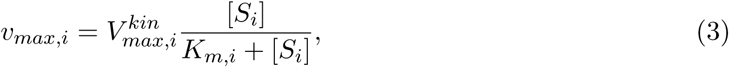

where 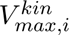 is the maximum kinetic uptake rate specified for substrate *i*, *K_m,i_* is the half-saturation concentration, and *v_max,i_*is the resulting concentration-dependent uptake limit, expressed in the standard FBA units of mmol gDW*^−^*^1^ h*^−^*^1^ (Supplementary Methods S1). Second, an additional mass-availability constraint limits uptake to the amount of substrate physically available within the local voxel during a single time-step (Supplementary Methods S3). The more restrictive of these two limits is applied to the corresponding exchange reaction. The resulting fluxes are then converted from dry-weight-normalised units (mmol gDW*^−^*^1^ h*^−^*^1^) to cell-level exchange rates (mmol min*^−^*^1^ per cell) and biomass growth using each agent’s instantaneous dry weight and volume (Supplementary Methods S2), providing a physically consistent mapping between intracellular metabolic fluxes, extracellular substrate exchange, and cellular growth; the resulting net export rate 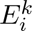 is used directly as the source or sink term for cell *k* in Eq. (2). Because per-cell exchange rates are small on the mmol min*^−^*^1^ scale, they are equivalently reported as fmol s*^−^*^1^ in some figures in the main text (1 mmol min*^−^*^1^ = 1.667 *×* 10^10^ fmol s*^−^*^1^); both notations refer to the same 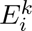 quantity. When biomass production is used as the optimisation objective, the resulting biomass flux is converted into a cell growth rate and used to update cell volume; other quantities derived from the FBA solution can also be coupled to PhysiCell phenotype and behavioural processes, including metabolism-driven motility and metabolism-dependent death. The complete derivation, dimensional scaling, unit conventions, validation, and symbol definitions are provided in Supplementary Methods S1–S5, and the full symbol glossary in Supplementary Table S12.

### Simulation scenarios

#### Genome-scale metabolic models

Two genome-scale metabolic models of *Escherichia coli* were used across the bacterial use cases presented in this work: the *E. coli* core metabolic model [36] and the genome-scale reconstruction iML1515 [37]. The syntrophic consortium combined models of *Clostridium beijerinckii* (iCB925) [38] and *Methanosarcina barkeri* Fusaro (iMG746) [39], while the tumour microenvironment use case employed an MCF7-specific human genome-scale reconstruction [29]. All models except the MCF7 model were obtained directly in SBML format; the MCF7 model was obtained in MATLAB/COBRA format and converted to SBML. Before simulation, models were minimally preprocessed to harmonise metabolite and exchange-reaction naming conventions where required, particularly for the two-species consortium, and to remove blocked reactions and gap metabolites, defined as metabolites that participate exclusively in blocked reactions [40], to reduce model size and improve computational efficiency; no other structural modifications were introduced. The resulting SBML models, together with the scripts used for model conversion and preprocessing, are publicly available at https://github.com/PhysiCelldFBA/dfba_sbml_models and through Zenodo [41]. A detailed description of the preprocessing workflow is provided in Supplementary Methods S6.

#### Mass conservation validation

A single *E. coli* cell, parameterised with the *E. coli* core model, was placed in a closed domain containing 10 mM glucose as the sole carbon source, with oxygen supplied without limitation and no further substrate input. The simulation was run until glucose exhaustion, and the resulting biomass trajectory was compared against the analytical prediction derived in Supplementary Methods S4. The validation was repeated using iML1515 in place of the core model to confirm that the result was independent of the specific metabolic reconstruction used. Complete simulation parameters and initial conditions are provided in Supplementary Methods S5.1.

#### Acetate switch

A well-mixed population of *E. coli* cells, parameterised with the *E. coli* core model, was simulated in a three-dimensional domain, with cells initially distributed randomly throughout the domain and glucose and oxygen supplied uniformly. The acetate exchange reaction was left unconstrained in the secretion direction, while its uptake bound was dynamically constrained by the local extracellular acetate concentration according to the kinetic and mass-availability rules described above; no additional regulatory logic was imposed on the glucose–acetate switch. At *t* = 3.5 h, extracellular oxygen was instantaneously reset to its initial concentration to represent re-aeration of the culture, as would occur upon shaking the flask, allowing the response to renewed oxygen availability and the resulting reactivation of aerobic acetate consumption to be evaluated as an in-silico representation of the acetate switch observed during aerobic *E. coli* growth. Complete simulation parameters and initial conditions are provided in Supplementary Methods S5.2.

#### Microbial colony

An expanding colony of *E. coli* cells, parameterised with the *E. coli* core model, was initialized as a three-dimensional colony with a radius of 10 *µ*m in a domain supplied with glucose and oxygen through Dirichlet boundary conditions and simulated for 10 h. The extracellular environment was initially supplied with 27.5 mM glucose and 25 mM oxygen, with glucose and oxygen maintained at these concentrations along the lateral domain boundaries. Cell growth, division, and mechanical displacement were governed by the standard PhysiCell cell-cycle and mechanics modules, with each cell’s growth rate determined independently by its own FBA solution. Complete simulation parameters and initial conditions are provided in Supplementary Methods S5.3.

#### Tumour microenvironment

A population of MCF7 breast cancer cells, parameterised with an MCF7-specific genome-scale reconstruction constrained by exometabolomic measurements, was arranged as a two-layer tissue section adjacent to a fixed blood vessel represented by a nutrient-supplying boundary. The vessel boundary supplied oxygen, glucose, and other extracellular metabolites at specified concentrations, while the corresponding extracellular fields were initially set to zero; the microenvironment was then evolved by diffusion until the metabolite concentration profiles stabilised, and the resulting spatial gradients were used as the initial extracellular conditions for the cellular simulation. Before spatial simulation, a phenotype phase plane analysis of the metabolic model was used to characterise the relationship between oxygen uptake, lactate secretion, and biomass production. This analysis revealed a broad region of equivalent or near-equivalent growth across multiple combinations of oxygen and lactate flux, indicating alternative optimal solutions (Supplementary Figure S7). The model was therefore operated near the transition between oxygen-replete and oxygen-limited growth regimes, where changes in local oxygen availability produce distinct proliferative, hypoxic, and necrotic phenotypes. Complete boundary conditions, diffusion parameters, and initial conditions are provided in Supplementary Methods S5.4.

#### Syntrophic consortium

A two-species community composed of the fermentative bacterium *Clostridium beijerinckii* (*i*CB925) and the methanogenic archaeon *Methanosarcina barkeri* Fusaro (*i*MG746) was seeded in a shared domain supplied with glucose and simulated for 50 h. The initial extracellular environment contained 1 mM glucose, 5 mM ammonium, 5 mM phosphate, and 5 mM sulfate, with CO_2_ set to 1 and methane, H_2_, and acetate initially absent; these concentrations were maintained at the domain boundaries. *C. beijerinckii* cells were supplied with glucose, with hydrogen and acetate production emerging from their local FBA solutions, while *M. barkeri* cells could consume hydrogen and CO_2_ through hydrogenotrophic methanogenesis or acetate through acetoclastic methanogenesis. Both methanogenic pathways were available in the metabolic model, and their relative utilisation was determined by local extracellular substrate availability and the resulting FBA solution rather than by a prescribed pathway-selection rule. Complete simulation parameters and initial conditions are provided in Supplementary Methods S5.5.

#### Metabolism-driven motility

A minimal model of metabolism-driven chemotaxis was implemented by combining PhysiCell’s built-in motility and chemotaxis modules with intracellular dFBA, parameterised with the *E. coli* core model. Migration speed was coupled to the ATP surplus available after basal maintenance, while a hysteretic phenotypic switch with distinct activation and deactivation thresholds on local glucose concentration governed the transition between motility- and growth-prioritising metabolic objectives. ATP maximisation was used for the motility-prioritising state and biomass maximisation for the growth-prioritising state. Two behavioural regimes were simulated using different glucose transport parameters, with *K_m,_*_Glc_ = 0.8 mM for the motility-dominated variant and *K_m,_*_Glc_ = 0.02 mM for the growth-recovery variant; the latter also used an increased ATP cost of maximum motility. Complete simulation parameters and initial conditions are provided in Supplementary Methods S5.6.

### Implementation

PhysiCelldFBA is implemented in C++ as an extension of PhysiCell v1.14.2, following the architectural design established by PhysiBoSS, in which the intracellular model is implemented as a concrete subclass of PhysiCell’s abstract intracellular model interface. This design allows PhysiCelldFBA to be attached to individual cell agents and updated alongside other intracellular models supported by the PhysiCell ecosystem without modifying the core PhysiCell simulation engine. Metabolic models are provided in SBML format and imported using the open-source library libSBML version 5.20.4 [42, 43]. Each cell agent maintains an independent metabolic model instance and computes an optimal flux distribution by solving the corresponding linear programme using the open-source simplex solver COIN-OR Clp version 1.17.10 [44] at every diffusion time-step, with the metabolic objective, extracellular substrate–exchange mappings, and growth parameters defined through the PhysiCell cell-definition configuration. Examples of dFBA configuration using XML and YAML files, together with the corresponding PhysiCell-Studio graphical interface, are shown in Supplementary Figures S2–S4. A summary of the documentation is provided in the Supplementary Note, while the full documentation, including examples and usage instructions, is available at https://physicelldfba.github.io/PhysiCelldFBA/.

### Data and Code Availability

The PhysiCelldFBA source code, example models, configuration files, and all simulation scenarios presented in this work are openly available at https://github.com/PhysiCelldFBA/PhysiCelldFBA/. All genome-scale metabolic models used in this study are available from the sources cited in the Simulation Scenarios section, and curated SBML versions of all models are publicly available on Zenodo [41]. No simulation output datasets are deposited, as all figures can be reproduced by executing the example scenarios included in the PhysiCelldFBA repository, and no new experimental datasets were generated. Any additional information required to reanalyse the data reported in this paper is available from the corresponding author upon request.

### Quantification and Statistical Analysis

No inferential statistical tests were performed in this study. All results presented are deterministic outputs of the PhysiCelldFBA simulation framework under the parameter sets specified in the Simulation Scenarios and Supplementary Methods S6. Validation of the mass-conservation coupling (Figure 1) was performed by comparing simulated biomass trajectories against analytical predictions derived from the initial glucose content of the domain and the biomass yield returned by the underlying genome-scale metabolic model; the relative error between simulated and predicted values is reported directly as a percentage. Spatial metabolite profiles in the tumour use case (Figure 4B) are shown as mean *±* standard deviation across all cells within 10 *µ*m distance bins. Cell-to-cell variability in metabolic fluxes is represented by shaded regions in the same figure. All figures were generated using Python 3.10.12 with Pandas 2.3.3, Matplotlib 3.10.9, and Seaborn 0.13.2.

## Author contributions

Conceptualisation: M.P., A.V. and P.M.; mathematical formulation: M.P. and M.R.; methodology and software: M.R., O.H.M. and M.P.; in silico experiments and simulations: M.R., O.H.M. and M.P.; results analysis: M.R., O.H.M. and M.P.; PhysiCell core software support and advising: R.H. and P.M.; platform test and compatibility: R.H. and M.R.; writing – original draft preparation: M.R., O.H.M. and M.P.; writing – review and editing: M.R., O.H.M., R.H., P.M., A.V. and M.P.; visualisation: M.R., O.H.M. and M.P.; supervision: M.P. and A.V. All authors have read and agreed to the published version of the manuscript.

## Ethics statement

This study is purely computational and did not involve human participants, human data, animal experimentation, or the collection of new biological samples. All genome-scale metabolic models used in this work were obtained from previously published, publicly available sources (see Simulation Scenarios and Data and Code Availability). No ethical approval was required for this study.

## Declaration of generative AI and AI-assisted technologies in the writing process

During the preparation of this work, the authors used LLMs to assist with language editing and manuscript organization. The authors reviewed and edited the generated content and take full responsibility for the content of the published article.

---

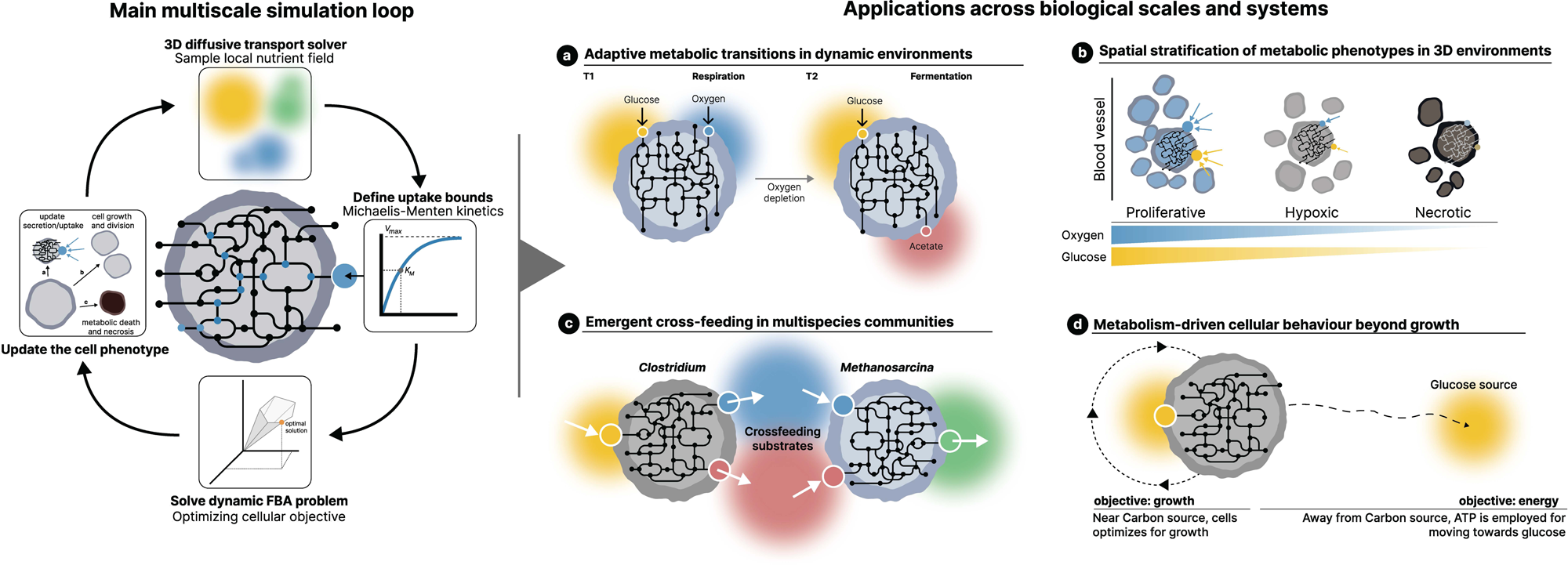

