## Supplementary Materials for "Integrating genome-scale metabolic models into multi-scale simulations"

### Supplementary Methods

#### S1. PhysiCelldFBA computational workflow

Figure S1 summarises the computational workflow implemented by PhysiCelldFBA. The framework extends the standard PhysiCell simulation loop by introducing a genome-scale metabolic optimisation step for each cell and coupling its outputs to the extracellular microenvironment and cell-level phenotype.

At each diffusion timestep, the following sequence is performed for each PhysiCelldFBA cell:

1. **Environmental sensing.** The cell samples the concentrations of extracellular substrates from its local BioFVM voxel.
2. **Metabolic constraint update.** Local substrate concentrations are used to update the bounds of the corresponding metabolic exchange reactions. Uptake is constrained by both concentration-dependent transport kinetics and the amount of substrate physically available within the local voxel.
3. **Metabolic optimisation.** The constrained genome-scale metabolic model is solved independently for the cell to obtain an optimal flux distribution.
4. **Metabolic-to-cellular coupling.** The resulting flux distribution provides metabolic information that can be coupled to multiple cell-level processes. Biomass production flux is converted into a cell-specific growth rate and used to update cell volume, while other metabolic fluxes can be used to control PhysiCell behavioural processes. For example, ATP production determines migration speed in the metabolism-driven motility scenario, while metabolic conditions can also trigger phenotype-dependent cell death.
5. **Metabolite exchange.** Exchange fluxes are converted from the dry-weight-normalised units used by the metabolic model to per-cell rates and passed to BioFVM, where they modify the local extracellular concentrations.
6. **Microenvironment and phenotype update.** BioFVM updates the extracellular reaction–diffusion system, while biomass production and other metabolic outputs update the corresponding PhysiCell cell states and behavioural processes.
7. **Iteration.** The updated extracellular concentrations and cell states provide the inputs for the next diffusion timestep.

This iterative coupling creates a feedback loop between intracellular metabolism, extracellular transport, and cell-level behaviour. Consequently, metabolic states can evolve in response to spatially and temporally varying environmental conditions, while metabolic activity can modify cellular growth, phenotype, motility, death, and the extracellular microenvironment. The detailed mathematical formulation of the exchange constraints, dimensional scaling, and local mass-conservation procedure is provided in Supplementary Methods S2–S3.

**dFBA configuration and model coupling** The dFBA behaviour of an agent is specified declaratively through a dedicated `<intracellular type="dfba">` block nested within the `<phenotype>` of a `<cell_definition>` in the PhysiCell configuration file (Supplementary Figure S2). PhysiCell defines cell types through independent `<cell_definition>` elements, each carrying its own phenotype specification (cycle, death, mechanics, motility, secretion, and so on). PhysiCelldFBA extends this mechanism so that the metabolic model becomes part of the cell type definition. Different `<cell_definition>` blocks can therefore reference different SBML models, or the same model with different transport or growth parameters, allowing heterogeneous metabolic populations, such as distinct species or distinct metabolic states of the same species, to coexist within a single simulation without modifying the underlying solver.

The `<growth_model>` block specifies how the selected objective reaction is coupled to the cellular phenotype. The `objective_reaction` entry identifies the FBA reaction used as the optimisation target, typically a biomass reaction but not necessarily so. When biomass production is the optimisation target, the resulting biomass flux is converted into a cell growth rate and used to update cell volume, while exchange fluxes defined in the `<exchange>` elements of the `<transport_model>` are used to update the extracellular environment. The `max_growth_rate` parameter sets an upper bound on biomass-associated growth, while `cell_density` and `reference_volume` are used to convert the predicted growth rate into cell-volume changes according to the procedure described in Supplementary Methods S2.

The metabolic solution can also be coupled to other PhysiCell phenotype and behavioural processes. In scenarios where biomass production is not the relevant phenotype, alternative objective reactions or metabolic fluxes can be used to drive cell behaviour, such as ATP-dependent motility, while metabolic conditions can also be used to trigger cell death. The corresponding scenario-specific configurations are described in Supplementary Methods S5.

#### S2. Biomass scaling and flux conversion

FBA exchange fluxes are expressed in mmol $\mathrm{gDW}^{-1}\,\mathrm{h}^{-1}$, whereas PhysiCell represents cells by their volume and BioFVM requires cell-level net export rates. PhysiCelldFBA therefore converts the fluxes returned by FBA using the instantaneous dry weight of each cell.

**Dry-weight estimation** PhysiCell stores the total cell volume $V_{cell}$ ($\mu m^3$) and the fluid fraction $\phi_{fluid}$. The corresponding solid volume is

$$V_{solid} = V_{cell}(1 - \phi_{fluid}). \tag{1}$$

Assuming a constant cellular dry-mass density $\rho_{cell}$ ($\mathrm{pg}\,\mu m^{-3}$), the dry weight of an individual cell is

$$m_{DW} = V_{solid}\,\rho_{cell}\,10^{-12}, \tag{2}$$

where the factor $10^{-12}$ converts picograms to grams.

**Scaling exchange fluxes** For substrate $i$, the FBA exchange flux $v_i$ is multiplied by the cell dry weight and converted from hours to minutes to obtain the cell-specific net export rate

$$E_i^k = \frac{v_i m_{DW}}{60}, \tag{3}$$

with units of $\mathrm{mmol\,min}^{-1}$ per cell. Positive and negative values of $E_i^k$ represent secretion and uptake, respectively, following the FBA exchange-flux sign convention. These rates provide the source or sink terms for the corresponding extracellular substrates in the BioFVM reaction–diffusion equation presented in the main text Methods.

For visualisation, cell-level export rates may be rescaled from $\mathrm{mmol\,min}^{-1}$ to femtomoles per minute or second.

**Growth-rate scaling** When biomass is used as the optimisation objective, the resulting biomass flux is interpreted as the specific growth rate $\mu$ ($\mathrm{h}^{-1}$). Because PhysiCell updates cell state variables on a minute-based timescale, the growth rate is converted to $\mathrm{min}^{-1}$ before updating cell volume. Assuming constant cellular dry-mass density, volumetric growth follows the biomass growth rate:

$$V(t + \Delta t) = V(t)\exp\left(\frac{\mu}{60}\Delta t\right), \tag{4}$$

where $\Delta t$ is the intracellular simulation timestep in minutes.

This formulation provides a consistent mapping between the biomass production predicted by the metabolic model and the geometric growth of individual PhysiCell agents.

#### S3. Local mass conservation algorithm

The kinetic uptake bound alone does not account for the finite amount of substrate physically present within a BioFVM voxel during a single timestep. An additional mass-availability constraint is therefore applied to each exchange reaction.

BioFVM voxel volumes are expressed in $\mu m^3$. To convert voxel volume to litres,

$$\alpha = 10^{-15}\ \mathrm{L}\,\mu m^{-3}. \tag{5}$$

For each exchangeable substrate $i$, the amount available in a voxel is

$$M_{voxel,i} = [S_i]\,V_{voxel}\,\alpha, \tag{6}$$

where $V_{voxel}$ is the voxel volume in $\mu m^3$.

The maximum uptake flux that can be sustained during the diffusion-synchronised timestep is then

$$v_{limit,i} = \frac{60\,M_{voxel,i}}{m_{DW}\,\Delta t_{intracellular}}, \tag{7}$$

where the factor 60 converts the timestep from minutes to hours. The resulting mass-availability limit is expressed in the same units as the kinetic uptake bound and can therefore be applied directly to the FBA exchange reaction.

The lower bound applied to the corresponding uptake exchange reaction is

$$v_{lb,i} = -\min\left(v_{max,i}, v_{limit,i}\right), \tag{8}$$

where $v_{max,i}$ is the kinetic uptake limit defined in the main text Methods. Thus, the exchange reaction is constrained by whichever limit is more restrictive. Because PhysiCelldFBA permits uptake from a given BioFVM voxel by only one cell at a time, this constraint ensures that the uptake applied during a single timestep cannot remove more substrate mass than is physically present in the voxel. Consequently, extracellular substrate concentrations remain non-negative after each update, independently of voxel size or uptake kinetics.

#### S4. Unit-test validation

The objective of this unit test is to verify that PhysiCelldFBA correctly couples extracellular substrate consumption, intracellular metabolic fluxes, biomass accumulation, and cell-based growth while conserving mass. Because the initial amount of substrate in the system is known, the maximum biomass that can be produced can be predicted analytically and directly compared with the simulation outcome. This provides a controlled benchmark for validating the dry-weight scaling procedure (Supplementary Methods S2) and the local mass-conservation algorithm (Supplementary Methods S3). The corresponding simulation results are presented in Figure 1 of the main text; this section details the analytical calculations underlying that comparison.

**Simulation setup** A single *E. coli* cell was placed in a closed environment containing an initial glucose concentration of $[Glc]_0 = 10$ mM ($0.01\ \mathrm{mol\,L^{-1}}$). The computational domain volume, converted to litres using the voxel-to-litre conversion factor introduced in Supplementary Methods S3 (Eq. (5)), was

$$V_{domain} = 4.096 \times 10^{-12}\ \mathrm{L}, \tag{9}$$

corresponding to an initial glucose mass of

$$M_{Glc} = [Glc]_0 \times V_{domain} \times MW_{Glc} = 7.38 \times 10^{-12}\ \mathrm{g}, \tag{10}$$

where $MW_{Glc} = 180.15\ \mathrm{g\,mol^{-1}}$ is the molecular weight of glucose.

**Analytical biomass prediction** The analytical prediction was derived directly from the same metabolic model used during the PhysiCelldFBA simulation, rather than from a fixed literature biomass yield. The *E. coli* metabolic reconstruction was solved independently using FBA under the same environmental conditions used in the simulation, and the biomass yield was calculated from the optimal biomass produced rate ($\mu$) and glucose uptake flux ($q_{Glc}$) as

$$Y_{X/S} = \frac{\mu}{q_{Glc}} \frac{1000}{MW_{Glc}} \quad \left(\mathrm{g\,DW\,(g\,Glc)^{-1}}\right), \tag{11}$$

where $q_{Glc}$ is expressed in mmol/gDW cell/hour and $\mu$ in $\mathrm{h^{-1}}$.
The theoretical biomass generated from the available glucose is therefore

$$\Delta M_X = Y_{X/S} \times M_{Glc}. \tag{12}$$

Because the simulation starts with a single founder cell, the analytical prediction reported in Figure 1 includes the biomass of the founder cell,

$$M_X^{theo} = M_{X,0} + \Delta M_X, \tag{13}$$

where $M_{X,0}$ is obtained from the PhysiCell reference cell volume, density, and solid fraction described in Supplementary Methods S2 (Eq. (2)).

**Simulation analysis** The total biomass at the end of the simulation was obtained from the final cell volumes using the same dry-weight scaling procedure described in Supplementary Methods S2. To ensure a consistent comparison with the analytical prediction, the founder cell biomass was included in both calculations.

To assess the robustness of the analytical prediction with respect to the choice of metabolic reconstruction, the same calculation was repeated using the genome-scale model iML1515 of *E. coli* [1] in place of the core model. The resulting biomass yield and theoretical biomass values were consistent with those obtained from the core reconstruction, indicating that the analytical benchmark is not specific to the core metabolic reconstruction and supporting the generality of the validation.

The validation was performed both with and without the non-growth-associated ATP maintenance (ATPM) constraint. Table S1 compares the analytical predictions with the corresponding PhysiCelldFBA simulations.

The analytical predictions and PhysiCelldFBA simulations agree to within 1% under both metabolic configurations. Because the simulation is performed in a closed system with a known initial glucose pool, this close agreement demonstrates that extracellular substrate consumption, intracellular flux balance calculations,

Table S1: Comparison between analytical predictions and PhysiCelldFBA simulations for the mass-conservation unit test.

| Condition | Analytical (pgDW) | Simulation (pgDW) | Relative error (%) |
|---|---|---|---|
| Without ATPM | 4.08 | 4.09 | 0.24 |
| With ATPM | 3.87 | 3.88 | 0.26 |

biomass accumulation, and agent-based growth remain quantitatively consistent while conserving mass throughout the simulation. These results validate the implementation of the dry-weight scaling procedure and support the correctness of the PhysiCelldFBA coupling between metabolism, substrate transport, and cell growth.

#### S5. Extended simulation parameterization

This section collects the parameter sets for the different PhysiCelldFBA examples reported in the main text that include: a minimal unit-test validation of the dFBA coupling; the *Escherichia coli* acetate switch; *E. coli* colony growth on the core metabolic model; A cancer-like tissue microenvironment; syntrophic cross-feeding between *Clostridium beijerinckii* and *Methanosarcina barkeri*; and the coupling of intracellular metabolism to *E. coli* motility and chemotaxis. Biological motivation, model selection, and interpretation of results are discussed in the corresponding subsections of the main text.

For each use case, a simulation-context table reports the computational domain and integration schedule, extracellular substrate initial concentration and boundary conditions, and the cellular and growth parameters that link FBA biomass flux to PhysiCell agent volume. Boundaries listed as **Closed (zero flux)** correspond to a disabled PhysiCell Dirichlet condition; fixed concentrations indicate an enabled Dirichlet boundary. Literature-based diffusion coefficients are summarised in Table , with scenario-specific modifications noted in Section S5.7. Intracellular transport kinetics for each metabolic model are collected in Section S5.7 (Tables S8, S9 and S10), and Section S5.8 compares the PhysiCelldFBA assignment of $K_m$, $V_{\max}$ and diffusion coefficients with similar simulation frameworks. Symbols are defined in Table S12.

##### S5.1. Unit test simulation scenario

This minimal *Escherichia coli* core-model scenario exchanges three metabolites: glucose, acetate and oxygen, to verify the dFBA coupling independently of any specific biological application [2]. Cell death was disabled so that the extracellular mass balance remains closed throughout the simulation. Domain, timesteps, extracellular conditions and cellular parameters are summarised in Table S2; diffusion coefficients follow Table . Transport kinetics follow Table S8, with the unit-test-specific adjustments described in Section S5.7.

Table S2: Simulation context for the dFBA unit test scenario.

| Parameter | Value | Unit | Description |
|---|---|---|---|
| Computational domain | $16 \times 16 \times 16$ | $\mu m^3$ | Simulation domain |
| Voxel size | $1.6 \times 1.6 \times 1.6$ | $\mu m^3$ | BioFVM mesh resolution |
| Simulation time | 4 | hours | Total simulated time |
| Diffusion timestep | 0.01 | min | BioFVM diffusion solver |
| Mechanics timestep | 0.1 | min | Cell mechanics |
| Phenotype timestep | 6 | min | Cell phenotype updates |
| Intracellular timestep | 0.01 | min | dFBA updates |
| *Extracellular substrate conditions* | | | |
| $CO_2$, initial concentration ($[CO_{2i}]_0$) | 0.0 | mM | Initial condition |
| $CO_2$, boundary condition | Closed (zero flux) | – | Dirichlet disabled (closed, zero flux) |
| Acetate, initial concentration ($[Ac_i]_0$) | 0.0 | mM | Initial condition |
| Acetate, boundary condition | Closed (zero flux) | – | Dirichlet disabled (closed, zero flux) |
| Glucose, initial concentration ($[Glc_i]_0$) | 10.0 | mM | Initial condition |
| Glucose, boundary condition | Closed (zero flux) | – | Dirichlet disabled (closed, zero flux) |
| Oxygen, initial concentration ($[O_{2i}]_0$) | Unlimited | mM | Initial condition |
| Oxygen, boundary condition | Closed (zero flux) | – | Dirichlet disabled (closed, zero flux) |
| *Cellular and growth parameters* | | | |
| Cellular dry-mass density ($\rho_{\text{cell}}$) | 1.04 | $g\,mL^{-1}$ | Biomass–volume coupling |
| Reference cell volume ($V_{\text{cell}}$) | 1.3 | $\mu m^3$ | Division / volume reference |
| Maximum specific growth rate ($\mu$) | 0.8 | $h^{-1}$ | Cap on FBA-linked growth |
| FBA biomass objective | `R_BIOMASS_Ecoli_core_w_GAM` | – | GEM objective reaction |
| Death model | Disabled | – | Closed mass-balance check |

##### S5.2. E. coli acetate switch simulation scenario

This *E. coli* core-model scenario reproduces overflow acetate secretion under oxygen-limiting growth on glucose and the subsequent re-consumption of acetate after glucose depletion [2, 3]. Simulation settings are summarised in Table S3, with diffusion coefficients from Table . Substrate-specific $K_m$ and $V_{\max}$ values (Table S8) were chosen so that the differing affinities for glucose, oxygen and acetate could capture the diauxic switch. The glucose Michaelis constant ($K_{m,\text{Glc}} = 0.02$ mM) is of the same order of magnitude as the value used in the original dynamic FBA study of mixed-substrate *E. coli* growth ($K_m = 0.015$ mM) [4]. Glucose and oxygen were assigned the highest uptake capacities, acetate an intermediate $V_{\max}$ supporting both overflow secretion and later re-consumption [3, 4], and $CO_2$ was treated as a freely exchanged by-product ($V_{\max} = 0$).

Table S3: Simulation context for the E. coli acetate switch scenario.

| Parameter | Value | Unit | Description |
|---|---|---|---|
| Computational domain | $700 \times 700 \times 3$ | $\mu m^3$ | Simulation domain |
| Voxel size | $1 \times 1 \times 1$ | $\mu m^3$ | BioFVM mesh resolution |
| Simulation time | 10 | h | Total simulated time |
| Diffusion timestep | 0.01 | min | BioFVM diffusion solver |
| Mechanics timestep | 0.1 | min | Cell mechanics |
| Phenotype timestep | 6 | min | Cell phenotype updates |
| Intracellular timestep | 0.01 | min | dFBA updates |
| *Extracellular substrate conditions* | | | |
| $CO_2$, initial concentration ($[CO_{2i}]_0$) | 0.0 | mM | Initial condition |
| $CO_2$, boundary condition | Closed (zero flux) | – | Dirichlet disabled (closed, zero flux) |
| Acetate, initial concentration ($[Ac_i]_0$) | 0.0 | mM | Initial condition |
| Acetate, boundary condition | Closed (zero flux) | – | Dirichlet disabled (closed, zero flux) |
| Glucose, initial concentration ($[Glc_i]_0$) | 8.0 | mM | Initial condition |
| Glucose, boundary condition | Closed (zero flux) | – | Dirichlet disabled (closed, zero flux) |
| Oxygen, initial concentration ($[O_{2i}]_0$) | 5 | mM | Initial condition |
| Oxygen, boundary condition | Closed (zero flux) | – | Dirichlet disabled (closed, zero flux) |
| *Cellular and growth parameters* | | | |
| Cellular dry-mass density ($\rho_{cell}$) | 1.04 | $g\,mL^{-1}$ | Biomass–volume coupling |
| Reference cell volume ($V_{cell}$) | 1.3 | $\mu m^3$ | Division / volume reference |
| Maximum specific growth rate ($\mu$) | 0.86 | $h^{-1}$ | Cap on FBA-linked growth |
| FBA biomass objective | `R_Biomass_Ecoli_core` | – | GEM objective reaction |

##### S5.3. E. coli colony growth simulation scenario

This colony-expansion scenario uses the *E. coli* core metabolic model [2]. Cells were initialised from a precomputed CSV file (`config/cells.csv`) with a nominal colony radius of 10 $\mu$m. Glucose and oxygen were held fixed at the domain boundaries (Dirichlet conditions of 27.5 and 25 mM, respectively) to mimic a nutrient-rich reservoir surrounding the expanding colony. Necrosis was enabled when the ATP maintenance flux (`R_ATPM`) exceeded a threshold of 8.39 mmol/gDW cell/hour. Simulation settings are summarised in Table S4, with diffusion coefficients from Table . Transport assumptions relative to the default core-model kinetics are detailed in the colony note of Section S5.7.

Table S4: Simulation context for the E. coli colony growth scenario.

| Parameter | Value | Unit | Description |
|---|---|---|---|
| Computational domain | $500 \times 500 \times 2$ | $\mu m^3$ | Simulation domain |
| Voxel size | $2 \times 2 \times 2$ | $\mu m^3$ | BioFVM mesh resolution |
| Simulation time | 10 | h | Total simulated time |
| Diffusion timestep | 0.001 | min | BioFVM diffusion solver |
| Mechanics timestep | 0.001 | min | Cell mechanics |
| Phenotype timestep | 0.001 | min | Cell phenotype updates |
| Intracellular timestep | 0.001 | min | dFBA updates |
| *Extracellular substrate conditions* | | | |
| $CO_2$, initial concentration ($[CO_{2i}]_0$) | 0.0 | mM | Initial condition |
| $CO_2$, boundary condition | 0.0 | mM | Dirichlet enabled (fixed concentration) |
| Acetate, initial concentration ($[Ac_i]_0$) | 0.0 | mM | Initial condition |
| Acetate, boundary condition | 0.0 | mM | Dirichlet enabled (fixed concentration) |
| Glucose, initial concentration ($[Glc_i]_0$) | 27.5 | mM | Initial condition |
| Glucose, boundary condition | 27.5 | mM | Dirichlet enabled (fixed concentration) |
| Lactate, initial concentration ($[Lac_i]_0$) | 0.0 | mM | Initial condition |
| Lactate, boundary condition | 0.0 | mM | Dirichlet enabled (fixed concentration) |
| Oxygen, initial concentration ($[O_{2i}]_0$) | 25.0 | mM | Initial condition |
| Oxygen, boundary condition | 25.0 | mM | Dirichlet enabled (fixed concentration) |
| *Cellular and growth parameters* | | | |
| Cellular dry-mass density ($\rho_{cell}$) | 1.04 | $g\,mL^{-1}$ | Biomass–volume coupling |
| Reference cell volume ($V_{cell}$) | 1.3 | $\mu m^3$ | Division / volume reference |
| Maximum specific growth rate ($\mu$) | 0.95 | $h^{-1}$ | Cap on FBA-linked growth |
| FBA biomass objective | `R_Biomass_Ecoli_core` | – | GEM objective reaction |
| Death model | necrosis | – | Triggered when `R_ATPM` exceeds 8.39 mmol/gDW cell/[illegible] |

##### S5.4. Cancer tissue simulation scenario

This MCF7 breast cancer scenario uses a trimmed Recon2.2 network. Nutrients are supplied at the $x_{min}$ boundary, and cells die by necrosis when the ATP-maintenance flux (`R_ATPM`) falls below a set threshold. Simulation settings are given in Table S5. Extracellular diffusion uses tissue-effective coefficients from the Cole hindrance relation for membrane-impermeable solutes (see Table ); diffusion values for $O_2$ and $CO_2$ are unchanged. Transport kinetics are listed in Table S9. Necrosis is triggered when `R_ATPM` cannot meet its lower bound (1.07 in model flux units). Reference cell volume and maximum growth rate convert FBA biomass flux to PhysiCell volume updates [5, 6].

Table S5: Simulation context for the MCF7 cancer tissue scenario (`PhysiCell_settings_wide_domain.xml`).

| Parameter | Value | Unit | Description |
|---|---|---|---|
| Computational domain | $360 \times 120 \times 30$ | $\mu m^3$ | Simulation domain |
| Voxel size | $15 \times 15 \times 15$ | $\mu m^3$ | BioFVM mesh resolution |
| Simulation time | 72 | h | Total simulated time (4320 min) |
| Diffusion timestep | 0.01 | min | BioFVM diffusion solver |
| Mechanics timestep | 0.1 | min | Cell mechanics |
| Phenotype timestep | 6 | min | Cell phenotype updates |
| Intracellular timestep | 0.01 | min | dFBA updates |
| *Extracellular substrate conditions (selected)* | | | |
| Glucose, initial concentration ($[Glc_i]_0$) | 0.0 | mM | Initial condition |
| Glucose, boundary ($x_{min}$) | 0.5 | mM | Dirichlet enabled |
| Glucose, boundary ($x_{max}$) | 0.2 | mM | Dirichlet enabled |
| Oxygen, initial concentration ($[O_{2i}]_0$) | 0.0 | mM | Initial condition |
| Oxygen, boundary ($x_{min}$) | 0.5 | mM | Dirichlet enabled |
| Oxygen, boundary ($x_{max}$) | 0.0 | mM | Dirichlet enabled |
| Lactate, initial concentration ($[Lac_i]_0$) | 0.0 | mM | Initial condition |
| Lactate, boundary ($x_{min}$) | Closed (zero flux) | – | Dirichlet disabled |
| Lactate, boundary ($x_{max}$) | 0.0 | mM | Dirichlet enabled |
| Gln and other amino acids, boundary ($x_{min}$) | 1.0 | mM | Dirichlet enabled (typical) |
| $CO_2$ / $NH_4^+$, boundary ($x_{min}$) | Closed (zero flux) | – | Dirichlet disabled |
| *Cellular and growth parameters* | | | |
| Cellular dry-mass density ($\rho_{cell}$) | 1.04 | $g\,mL^{-1}$ | Biomass–volume coupling |
| Reference cell volume ($V_{cell}$) | 2494 | $\mu m^3$ | Division / volume reference |
| Maximum specific growth rate ($\mu$) | 0.026803 | $h^{-1}$ | Cap on FBA-linked growth |
| FBA biomass objective | `R_biomass_reaction` | – | GEM objective reaction |
| Death model | necrosis | – | `R_ATPM` threshold 1.07; rate increment $1.67 \times 10^{-7}$ |

#### S5.5. Cross-feeding simulation scenario

This syntrophic co-culture combines *Clostridium beijerinckii* (CB) and *Methanosarcina barkeri* (MB) using the iCB925 and iMG746 reconstructions, respectively [7, 8]. CB ferments glucose and releases $H_2$, $CO_2$ and acetate, which MB consumes to produce methane. Simulation settings for both organisms are summarised in Table S6, with diffusion coefficients from Table . Transport kinetics are given separately for each species in Table S10; the $H_2$ Michaelis-constant adjustment for *M. barkeri* is discussed in Section S5.7.

Table S6: Simulation context for the microbial cross-feeding scenario.

| Parameter | Value | Unit | Description |
|---|---|---|---|
| Computational domain | $200 \times 200 \times 1.5$ | $\mu m^3$ | Simulation domain |
| Voxel size | $0.8 \times 0.8 \times 0.8$ | $\mu m^3$ | BioFVM mesh resolution |
| Simulation time | 50 | h | Total simulated time |
| Diffusion timestep | 0.01 | min | BioFVM diffusion solver |
| Mechanics timestep | 0.1 | min | Cell mechanics |
| Phenotype timestep | 6 | min | Cell phenotype updates |
| Intracellular timestep | 0.01 | min | dFBA updates |
| *Extracellular substrate conditions* | | | |
| $CO_2$, initial concentration ($[CO_{2i}]_0$) | 1 | mM | Initial condition |
| $CO_2$, boundary condition | 1 | mM | Dirichlet enabled (fixed concentration) |
| $H_2$, initial concentration ($[H_{2i}]_0$) | 0 | mM | Initial condition |
| $H_2$, boundary condition | 0 | mM | Dirichlet enabled (fixed concentration) |
| $NH_4^+$, initial concentration ($[NH4_i]_0$) | 5 | mM | Initial condition |
| $NH_4^+$, boundary condition | 5 | mM | Dirichlet enabled (fixed concentration) |
| $P_i$, initial concentration ($[Pi_i]_0$) | 5 | mM | Initial condition |
| $P_i$, boundary condition | 5 | mM | Dirichlet enabled (fixed concentration) |
| $SO_4^{2-}$, initial concentration ($[SO4_i]_0$) | 5 | mM | Initial condition |
| $SO_4^{2-}$, boundary condition | 5 | mM | Dirichlet enabled (fixed concentration) |
| Acetate, initial concentration ($[Ac_i]_0$) | 0 | mM | Initial condition |
| Acetate, boundary condition | 0 | mM | Dirichlet enabled (fixed concentration) |
| Glucose, initial concentration ($[Glc_i]_0$) | 1 | mM | Initial condition |
| Glucose, boundary condition | 1 | mM | Dirichlet enabled (fixed concentration) |
| Methane, initial concentration ($[CH_{4i}]_0$) | 0.0 | mM | Initial condition |
| Methane, boundary condition | 0.0 | mM | Dirichlet enabled (fixed concentration) |
| *Cellular and growth parameters (C. beijerinckii)* | | | |
| Cellular dry-mass density ($\rho_{cell}$) | 1.04 | $g\,mL^{-1}$ | Biomass–volume coupling |
| Reference cell volume ($V_{cell}$) | 1.3 | $\mu m^3$ | Division / volume reference |
| Maximum specific growth rate ($\mu$) | 0.4 | $h^{-1}$ | Cap on FBA-linked growth |
| FBA biomass objective | `R_biomass` | – | GEM objective reaction |
| *Cellular and growth parameters (M. barkeri)* | | | |
| Cellular dry-mass density ($\rho_{cell}$) | 1.04 | $g\,mL^{-1}$ | Biomass–volume coupling |
| Reference cell volume ($V_{cell}$) | 1.8 | $\mu m^3$ | Division / volume reference |
| Maximum specific growth rate ($\mu$) | 0.12 | $h^{-1}$ | Cap on FBA-linked growth |
| FBA biomass objective | `R_Mb_biomass_65` | – | GEM objective reaction |

##### S5.6. Coupling intracellular metabolism to cell motility

This *E. coli* core-model scenario combines PhysiCell motility and chemotaxis with intracellular dFBA [2]. Migration speed is set from the ATP surplus available after basal maintenance, and a hysteretic phenotypic switch governs the transition between motility and biomass optimisation as a function of local glucose availability (see Materials and Methods). Cell division was enabled so that growth recovery could lead to proliferation once the metabolic objective switched from motility to biomass production. Simulation settings are summarised in Table S7, with diffusion coefficients from Table . Two behavioural regimes reported in the main text differ primarily in the glucose Michaelis constant (Table S8): the motility-dominated variant uses $K_{m,\mathrm{Glc}} = 0.8$ mM, yielding sufficient ATP for directed migration but insufficient glucose flux to trigger the return to biomass optimisation; the growth-recovery variant restores $K_{m,\mathrm{Glc}} = 0.02$ mM together with an increased ATP cost of maximum motility, allowing cells to recover volumetric growth once local glucose exceeds the upper switching threshold. Oxygen, acetate and $CO_2$ transport parameters are shared between variants.

Table S7: Simulation context for the metabolic-driven motility scenario.

| Parameter | Value | Unit | Description |
|---|---|---|---|
| Computational domain | 300 × 100 × 10 | $\mu m^3$ | Simulation domain |
| Voxel size | 5 × 5 × 10 | $\mu m^3$ | BioFVM mesh resolution |
| Simulation time | 10 | h | Total simulated time |
| Diffusion timestep | 0.01 | min | BioFVM diffusion solver |
| Mechanics timestep | 0.1 | min | Cell mechanics |
| Phenotype timestep | 6 | min | Cell phenotype updates |
| Intracellular timestep | 0.01 | min | dFBA updates |
| *Extracellular substrate conditions* | | | |
| $CO_2$, initial concentration ($[CO_{2i}]_0$) | 0.0 | mM | Initial condition |
| $CO_2$, boundary condition | Closed (zero flux) | – | Dirichlet disabled (closed, zero flux) |
| Acetate, initial concentration ($[Ac_i]_0$) | 0.0 | mM | Initial condition |
| Acetate, boundary condition | Closed (zero flux) | – | Dirichlet disabled (closed, zero flux) |
| Glucose, initial concentration ($[Glc_i]_0$) | 0.0 | mM | Initial condition |
| Glucose, boundary condition | 0.0 | mM | Dirichlet enabled (fixed concentration) |
| Oxygen, initial concentration ($[O_{2i}]_0$) | 4 | mM | Initial condition |
| Oxygen, boundary condition | 8 | mM | Dirichlet enabled (fixed concentration) |
| *Cellular and growth parameters* | | | |
| Cellular dry-mass density ($\rho_{cell}$) | 1.04 | $g\,mL^{-1}$ | Biomass–volume coupling |
| Reference cell volume ($V_{cell}$) | 10 | $\mu m^3$ | Division / volume reference |
| Maximum specific growth rate ($\mu$) | 0.86 | $h^{-1}$ | Cap on FBA-linked growth |
| FBA biomass objective | `R_Biomass_Ecoli_core` | – | GEM objective reaction |

##### S5.7. Organism metabolic and cellular parameters

This section collects the intracellular transport and extracellular diffusion parameters shared across Sections S5.1–S5.6. Default *E. coli* core-model kinetics are listed in Table S8, with scenario-specific modifications described in the accompanying notes. Diffusion coefficients are summarised in Table , and transport kinetics for the MCF7 and cross-feeding scenarios appear in Tables S9 and S10. Cellular and growth parameters are reported in the simulation-c ontext tables of Sections S5.1–S5.6. Symbols follow Table S12.

The **Reference** column cites the primary source for each non-zero $K_m$ or $V_{\max}^{\text{kin}}$. Where published dynamic-FBA calibrations or tumour-metabolism studies report affinities or uptake capacities, these are cited directly. Genome-scale reconstructions specify exchange-reaction bounds but not extracellular Michaelis–Menten parameters; in those cases the corresponding reconstruction is cited because uptake kinetics were assigned to bound the listed SBML exchange reactions consistently with each model's flux limits. Maximum uptake rates were scaled to remain compatible with exchange-reaction upper bounds while preserving literature $K_m$ order of magnitude where available. The suffix ", scenario-adjusted" marks explicit scenario overrides. The Reference column is left empty when $V_{\max}^{\text{kin}} = 0$ (metabolites secreted without re-uptake).

***Escherichia coli* core metabolic model** Table S8 lists default *E. coli* core-model transport kinetics (Sections S5.1, S5.2, S5.6). Two use cases depart from these defaults:

- **Unit test (S5.1).** $V_{\max,\text{Ac}}^{\text{kin}}$ was set to zero so that acetate is treated as a secreted by-product with no re-uptake, simplifying closed mass-balance checks of the dFBA–transport coupling.
- **Metabolic motility (S5.6).** The motility-dominated behavioural regime uses $K_{m,\text{Glc}} = 0.8$ mM rather than the default 0.02 mM, yielding sufficient ATP for directed migration but limiting glucose flux so that cells remain in the motility-optimised state; the growth-recovery variant restores $K_{m,\text{Glc}} = 0.02$ mM (see Section S5.6).

Colony growth (Section S5.3) applies the transport deviations summarised in **Note (colony)** below the table.

Table S8: Default *Escherichia coli* core-model transport kinetics for the PhysiCelldFBA scenarios (Sections S5.1, S5.2, S5.6). Scenario-specific overrides are described in the text above; colony deviations (Section S5.3) are given in **Note (colony)**. The Reference column cites the reconstruction or literature calibration for each parameter (non-zero values only).

| Symbol | Description | Value | Unit | Reference |
|---|---|---|---|---|
| $K_{m,\mathrm{Glc}}$ | Half-saturation constant for Glucose | 0.02 | mM | [4] |
| $V_{\mathrm{max,Glc}}^{\mathrm{kin}}$ | Maximum kinetic uptake rate constant for Glucose | 8.0 | mmol/gDW cell/hour | [4] |
| $K_{m,\mathrm{Ac}}$ | Half-saturation constant for Acetate | 1 | mM | [3] |
| $V_{\mathrm{max,Ac}}^{\mathrm{kin}}$ | Maximum kinetic uptake rate constant for Acetate | 5.0 | mmol/gDW cell/hour | [3, 4] |
| $K_{m,\mathrm{O_2}}$ | Half-saturation constant for Oxygen | 0.002 | mM | [4] |
| $V_{\mathrm{max,O_2}}^{\mathrm{kin}}$ | Maximum kinetic uptake rate constant for Oxygen | 18 | mmol/gDW cell/hour | [4] |
| $K_{m,\mathrm{CO_2}}$ | Half-saturation constant for $CO_2$ | 1 | mM | [2] |
| $V_{\mathrm{max,CO_2}}^{\mathrm{kin}}$ | Maximum kinetic uptake rate constant for $CO_2$ | 0.0 | mmol/gDW cell/hour | – |

**Notes.** Intracellular metabolism was represented using the *Escherichia coli* core metabolic model [2]. The reconstruction specifies exchange-reaction bounds but not extracellular concentrations or transporter kinetics; these phenomenological parameters were assigned in PhysiCelldFBA to translate local substrate concentrations into dynamic FBA uptake bounds. Values differ between scenarios: neutral reference constants ($K_m = V_{\mathrm{max}} = 1$) in the unit test; substrate-specific affinities and capacities in the acetate-switch and motility simulations to capture diauxic overflow metabolism and chemotactic gradients, respectively. Cell density was set to 1.04 g mL$^{-1}$; reference cell volume and maximum growth rate convert FBA biomass flux to PhysiCell volume updates.

**Note (colony).** Colony growth (Section S5.3) uses the *E. coli* core metabolic model [2]. Uptake parameters were assigned as deviations from the default core-model values in Table S8: Michaelis constants were set uniformly to $K_m = 0.01$ mM for all exchanged metabolites (versus substrate-specific core values); maximum kinetic uptake rates were $V_{\mathrm{max,Glc}}^{\mathrm{kin}} = 10.0$, $V_{\mathrm{max,Ac}}^{\mathrm{kin}} = 10.0$, and $V_{\mathrm{max,Lac}}^{\mathrm{kin}} = 10.0$ mmol/gDW cell/hour for glucose, acetate, and lactate (with lactate exchange added), $V_{\mathrm{max,O_2}}^{\mathrm{kin}} = 20.0$ mmol/gDW cell/hour for oxygen, and $V_{\mathrm{max,CO_2}}^{\mathrm{kin}} = 10.0$ mmol/gDW cell/hour for $CO_2$ (core default $V_{\mathrm{max,CO_2}}^{\mathrm{kin}} = 0$). Maximum uptake rates were scaled according to metabolic role (elevated for oxygen, uniform for carbon substrates) to limit the number of free parameters while allowing local depletion in the colony interior to progressively limit growth [4, 9]. Cell density was set to 1.04 g mL$^{-1}$. Necrosis was enabled when the ATP maintenance flux (`R_ATPM`) exceeded a threshold of 8.39 mmol gDW$^{-1}$ h$^{-1}$.

***MCF7* cancer cell line** Table S9 lists Michaelis–Menten transport parameters for the MCF7 Recon2.2 cancer scenario (Section S5.4), as configured in `PhysiCell_settings_wide_domain.xml`. Glucose, glutamine, lactate, and oxygen provide the primary carbon/redox coupling to the GEM; additional amino acids and micronutrients are assigned uniform supporting kinetics ($K_m = 1$ mM, $V_{\mathrm{max}}^{\mathrm{kin}} = 0.02$ mmol/gDW cell/hour, except glutamine at $V_{\mathrm{max,Gln}}^{\mathrm{kin}} = 0.1$).

Table S9: *MCF7* breast cancer cell line (Recon2.2): transport kinetics for the cancer scenario (Section S5.4). Units match the PhysiCell XML exchange block. Remaining amino-acid exchanges (Arg, Asn, Asp, Glu, Gly, Ile, Leu, Lys, Orn, Phe, Ser, Thr, Trp, Tyr, Val, Cho, $NH_4^+$, $P_i$) use $K_m = 1$ mM and $V_{\mathrm{max}}^{\mathrm{kin}} = 0.02$ mmol/gDW cell/hour unless noted.

| Symbol | Description | Value | Unit | Reference |
|---|---|---|---|---|
| $K_{m,\mathrm{Glc}}$ | Half-saturation constant for Glucose | 5.0 | mM | [5, 6], scenario-adjusted |
| $V_{\mathrm{max,Glc}}^{\mathrm{kin}}$ | Maximum kinetic uptake rate for Glucose | 1.0 | mmol/gDW cell/hour | [5], scenario-adjusted |
| $K_{m,\mathrm{Gln}}$ | Half-saturation constant for Glutamine | 1.0 | mM | [5, 6] |
| $V_{\mathrm{max,Gln}}^{\mathrm{kin}}$ | Maximum kinetic uptake rate for Glutamine | 0.1 | mmol/gDW cell/hour | [5] |
| $K_{m,\mathrm{Lac}}$ | Half-saturation constant for Lactate | 1.0 | mM | [5, 6] |
| $V_{\mathrm{max,Lac}}^{\mathrm{kin}}$ | Maximum kinetic uptake rate for Lactate | 0.02 | mmol/gDW cell/hour | [5], scenario-adjusted |
| $K_{m,\mathrm{O_2}}$ | Half-saturation constant for Oxygen | 0.5 | mM | [5, 6], scenario-adjusted |
| $V_{\mathrm{max,O_2}}^{\mathrm{kin}}$ | Maximum kinetic uptake rate for Oxygen | 1.0 | mmol/gDW cell/hour | [5], scenario-adjusted |
| $K_{m,\mathrm{CO_2}}$ | Half-saturation constant for $CO_2$ | 1.0 | mM | [5] |
| $V_{\mathrm{max,CO_2}}^{\mathrm{kin}}$ | Maximum kinetic uptake rate for $CO_2$ | 0.02 | mmol/gDW cell/hour | [5] |

**Notes.** The MCF7 cancer cell line was modelled with a trimmed Recon2.2 network [5]. A necrosis death model removes metabolically compromised cells when the ATP-maintenance reaction `R_ATPM` cannot satisfy its lower bound (1.07 model flux units; death-rate increment $1.67\times10^{-7}$). Reference cell density $\rho_{\mathrm{cell}} = 1.04$ g mL$^{-1}$, reference volume $V_{\mathrm{cell}} = 2494$ $\mu$m$^3$, and maximum growth rate $\mu = 0.026803$ h$^{-1}$ link FBA biomass flux to PhysiCell volume updates.

**Cross-feeding community GEMs** Table S10 lists transport kinetics for the syntrophic cross-feeding scenario (Section S5.5). Unlike the *E. coli* core and MCF7 scenarios, iCB925 and iMG746 do not report substrate-specific transporter kinetics; all uptake parameters were assigned from each model's exchange-reaction structure (finite $V_{\max}^{\mathrm{kin}}$ for consumed metabolites, $V_{\max}^{\mathrm{kin}} = 0$ for secreted by-products). Parameters are specified separately for each organism because the two populations share extracellular metabolite pools: consumed substrates receive finite uptake capacities, whereas secreted metabolites are assigned $V_{\max}^{\mathrm{kin}} = 0$ so that fermentation products released by *C. beijerinckii* accumulate and support methanogenesis in *M. barkeri*. For *M. barkeri*, $K_{m,\mathrm{H_2}}$ was raised from 0.0001 mM (near-saturating in earlier tests) to 0.05 mM, within the phenomenological range assigned for iMG746 (0.01–0.1 mM; [8]), to soften uptake coupling and better balance $H_2$ production with consumption.

Table S10: Transport kinetics for the cross-feeding scenario (Section S5.5): *Clostridium beijerinckii* (iCB925) and *Methanosarcina barkeri* (iMG746). Reference: reconstruction or literature calibration (non-zero parameters only).

| Symbol | Description | Value | Unit | Reference |
|---|---|---|---|---|
| *Clostridium beijerinckii* iCB925 (Section S5.5) | | | | |
| $K_{m,\mathrm{Glc}}$ | Half-saturation constant for Glucose | 1 | mM | [7] |
| $V_{\max,\mathrm{Glc}}^{\mathrm{kin}}$ | Maximum kinetic uptake rate constant for Glucose | 7.5 | mmol/gDW cell/hour | [7] |
| $K_{m,\mathrm{H_2}}$ | Half-saturation constant for $H_2$ | 0.1 | mM | [7] |
| $V_{\max,\mathrm{H_2}}^{\mathrm{kin}}$ | Maximum kinetic uptake rate constant for $H_2$ | 0.0 | mmol/gDW cell/hour | – |
| $K_{m,\mathrm{NH4}}$ | Half-saturation constant for $NH_4^+$ | 0.001 | mM | [7] |
| $V_{\max,\mathrm{NH4}}^{\mathrm{kin}}$ | Maximum kinetic uptake rate constant for $NH_4^+$ | 15.0 | mmol/gDW cell/hour | [7] |
| $K_{m,\mathrm{Pi}}$ | Half-saturation constant for $P_i$ | 0.001 | mM | [7] |
| $V_{\max,\mathrm{Pi}}^{\mathrm{kin}}$ | Maximum kinetic uptake rate constant for $P_i$ | 15.0 | mmol/gDW cell/hour | [7] |
| $K_{m,\mathrm{SO4}}$ | Half-saturation constant for $SO_4^{2-}$ | 0.001 | mM | [7] |
| $V_{\max,\mathrm{SO4}}^{\mathrm{kin}}$ | Maximum kinetic uptake rate constant for $SO_4^{2-}$ | 15.0 | mmol/gDW cell/hour | [7] |
| $K_{m,\mathrm{Ac}}$ | Half-saturation constant for Acetate | 0.1 | mM | [7] |
| $V_{\max,\mathrm{Ac}}^{\mathrm{kin}}$ | Maximum kinetic uptake rate constant for Acetate | 0.0 | mmol/gDW cell/hour | – |
| $K_{m,\mathrm{CO_2}}$ | Half-saturation constant for $CO_2$ | 0.1 | mM | [7] |
| $V_{\max,\mathrm{CO_2}}^{\mathrm{kin}}$ | Maximum kinetic uptake rate constant for $CO_2$ | 0.0 | mmol/gDW cell/hour | – |
| *Methanosarcina barkeri* iMG746 (Section S5.5) | | | | |
| $K_{m,\mathrm{CO_2}}$ | Half-saturation constant for $CO_2$ | 0.0001 | mM | [8] |
| $V_{\max,\mathrm{CO_2}}^{\mathrm{kin}}$ | Maximum kinetic uptake rate constant for $CO_2$ | 100.0 | mmol/gDW cell/hour | [8] |
| $K_{m,\mathrm{H_2}}$ | Half-saturation constant for $H_2$ | 0.05 | mM | [8], scenario-adjusted |
| $V_{\max,\mathrm{H_2}}^{\mathrm{kin}}$ | Maximum kinetic uptake rate constant for $H_2$ | 20.0 | mmol/gDW cell/hour | [8] |
| $K_{m,\mathrm{CH_4}}$ | Half-saturation constant for Methane | 0.0001 | mM | [8] |
| $V_{\max,\mathrm{CH_4}}^{\mathrm{kin}}$ | Maximum kinetic uptake rate constant for Methane | 0.0 | mmol/gDW cell/hour | – |
| $K_{m,\mathrm{NH4}}$ | Half-saturation constant for $NH_4^+$ | 0.0001 | mM | [8] |
| $V_{\max,\mathrm{NH4}}^{\mathrm{kin}}$ | Maximum kinetic uptake rate constant for $NH_4^+$ | 10.0 | mmol/gDW cell/hour | [8] |
| $K_{m,\mathrm{Pi}}$ | Half-saturation constant for $P_i$ | 0.0001 | mM | [8] |
| $V_{\max,\mathrm{Pi}}^{\mathrm{kin}}$ | Maximum kinetic uptake rate constant for $P_i$ | 10 | mmol/gDW cell/hour | [8] |
| $K_{m,\mathrm{Ac}}$ | Half-saturation constant for Acetate | 0.0001 | mM | [8] |
| $V_{\max,\mathrm{Ac}}^{\mathrm{kin}}$ | Maximum kinetic uptake rate constant for Acetate | 3.0 | mmol/gDW cell/hour | [8] |

**Notes (C. beijerinckii).** *Clostridium beijerinckii* metabolism was represented with the iCB925 genome-scale model [7]. Michaelis constants and maximum uptake rates were assigned according to the direction of each exchange: finite uptake capacities for consumed substrates (glucose, ammonium, phosphate, sulfate) and zero uptake capacity for secreted by-products ($H_2$, acetate, $CO_2$), so that fermentation products become available to the partner organism. Cell density, reference volume, and maximum growth rate provide the biomass–volume coupling described in the main text.

**Notes (M. barkeri).** *Methanosarcina barkeri* metabolism was represented with the iMG746 reconstruction [8]. Transport parameters were set so that $H_2$, $CO_2$, and acetate, which is released by *C. beijerinckii*, support methanogenesis, while methane was assigned zero uptake capacity. Mineral nutrients ($NH_4^+$, $P_i$) were maintained at exchange capacities consistent with the GEM bounds.

**Extracellular substrate diffusion coefficients** Table  lists reference diffusion coefficients for each exchanged metabolite. Each **Reference** entry cites the primary measurement or tabulation for that metabolite: the BioFVM default for $O_2$ [10]; micro-electrode biofilm measurements and the BioFVM default for glucose [10, 11]; diffusion measurements for acetate [12]; Taylor-dispersion gas-in-water measurements for $CO_2$ [13], $H_2$ [14], and $CH_4$ [15]; Taylor-dispersion measurements for lactate [16]; aqueous ion diffusivities for $NH_4^+$ [17]; early-diagenesis phosphorus transport [18]; and sulfate tracer diffusion [19]. These values are consistent with those adopted in spatial reaction–diffusion colony models [9] where applicable. The unit test (S5.1), acetate switch (S5.2), metabolic motility (S5.6), and cross-feeding (S5.5) adopt the literature coefficients without modification. Colony growth (S5.3) uses literature coefficients assigned from the same primary sources ($D_{\mathrm{O_2}} = 120000$, $D_{\mathrm{Glc}} = 35000$, $D_{\mathrm{Ac}} = 50000$, $D_{\mathrm{Lac}} = 50000$, $D_{\mathrm{CO_2}} = 100000$ $\mu\mathrm{m}^2$/min).

One scenario departs from the reference diffusion coefficients: The cancer tissue simulation uses *tissue-effective* diffusivities for membrane-impermeable solutes, obtained from the Cole volume-fraction hindrance relation [9].

- **Cancer tissue (S5.4).** The cancer tissue domain is a densely packed multicellular aggregate (15 $\mu$m voxels; cell centres on a lattice with packing fraction $\rho \approx 0.74$). For membrane-impermeable solutes such as glucose, extracellular diffusion occurs in the interstitial fluid and is hindered by the cell volume fraction. Following Cole [9],

$$D_{\text{eff}} = \frac{1-\rho}{1+\rho/2} D_{\text{water}}, \tag{14}$$

with aqueous glucose $D_{\text{water,Glc}} = 46800$ $\mu\text{m}^2/\text{min}$ (Table ) yielding $D_{\text{eff,Glc}} = 8921$ $\mu\text{m}^2/\text{min}$. All other membrane-impermeable extracellular metabolites in this scenario (amino acids, lactate, phosphate, ammonium, choline, etc.) were assigned the *same* tissue-effective diffusivity $D = D_{\text{eff,Glc}} = 8921$ $\mu\text{m}^2/\text{min}$, so that differential extracellular transport does not introduce spurious substrate-specific length scales beyond those set by uptake kinetics. Membrane-permeable gases were left at aqueous values: $D_{O_2} = 100000$ $\mu\text{m}^2/\text{min}$ [10] and $D_{CO_2} = 115000$ $\mu\text{m}^2/\text{min}$ [13], because Cole-type interstitial hindrance does not apply when solute crosses cell membranes freely.

Table S11: Literature substrate diffusion coefficients in water at 37 °C (reference tabulation). For the cancer scenario (S5.4), membrane-impermeable solutes use the Cole tissue-effective value $D_{\text{eff,Glc}} = 8921$ $\mu\text{m}^2/\text{min}$ (and the same $D$ for all other impermeable metabolites), while $O_2$ and $CO_2$ retain aqueous diffusivities; see the text above. Initial and boundary concentrations are given in the simulation-context tables for each scenario (Sections S5.1–S5.6).

| Metabolite | Symbol | Description | Value | Unit | Reference |
|---|---|---|---|---|---|
| Oxygen | $D_{O_2}$ | Diffusion coefficient for Oxygen | 100000 | $\mu\text{m}^2/\text{min}$ | [10] |
| Glucose | $D_{\text{Glc}}$ | Diffusion coefficient for Glucose | 46800 | $\mu\text{m}^2/\text{min}$ | [10, 11] |
| Acetate | $D_{\text{Ac}}$ | Diffusion coefficient for Acetate | 72000 | $\mu\text{m}^2/\text{min}$ | [12] |
| $CO_2$ | $D_{CO_2}$ | Diffusion coefficient for $CO_2$ | 115000 | $\mu\text{m}^2/\text{min}$ | [13] |
| Lactate | $D_{\text{Lac}}$ | Diffusion coefficient for Lactate | 60000 | $\mu\text{m}^2/\text{min}$ | [16] |
| Methane | $D_{CH_4}$ | Diffusion coefficient for Methane | 90000 | $\mu\text{m}^2/\text{min}$ | [15] |
| $H_2$ | $D_{H_2}$ | Diffusion coefficient for $H_2$ | 270000 | $\mu\text{m}^2/\text{min}$ | [14] |
| $NH_4^+$ | $D_{\text{NH4}}$ | Diffusion coefficient for $NH_4^+$ | 118000 | $\mu\text{m}^2/\text{min}$ | [17] |
| $P_i$ | $D_{\text{Pi}}$ | Diffusion coefficient for $P_i$ | 53000 | $\mu\text{m}^2/\text{min}$ | [18] |
| $SO_4^{2-}$ | $D_{\text{SO4}}$ | Diffusion coefficient for $SO_4^{2-}$ | 64000 | $\mu\text{m}^2/\text{min}$ | [19] |

##### Comparison of parametrisation with existing spatial dFBA frameworks

Spatial dynamic-FBA frameworks differ in how they assign the Michaelis–Menten uptake parameters ($K_m$, $V_{\max}$) and extracellular diffusion coefficients ($D$) that couple metabolite fields to intracellular FBA. Here we contrast the PhysiCelldFBA parametrisation with two widely used alternatives, COMETS [20] and BacArena [21].

**Diffusion coefficients.** COMETS assigns a single metabolite diffusion coefficient to all substrates ($D = 5 \times 10^{-6}$ $\text{cm}^2/\text{s}$, based on sugar diffusion in water [22]; $\approx 3.0 \times 10^4$ $\mu\text{m}^2/\text{min}$). BacArena uses a default `difspeed` of 0.02412 $\text{cm}^2/\text{h}$ ($\approx 6.7 \times 10^{-6}$ $\text{cm}^2/\text{s}$, corresponding to aqueous glucose) when substances are added to the arena, with the option to set $D$ independently for each metabolite. PhysiCelldFBA assigns metabolite-specific diffusion coefficients based on literature (Table ), and applies the Cole tissue-effective correction only in the cancer tissue scenario (Section S5.4), where membrane-impermeable solutes share $D_{\text{eff,Glc}}$ while $O_2/CO_2$ are assumed to be aqueous. The three approaches are consistent in order of magnitude for small solutes ($D_{\text{Glc}} = 46800$ $\mu\text{m}^2/\text{min}$ in Table ).

**Michaelis constants ($K_m$).** COMETS applies a common $K_m$ to all exchanged metabolites in its uptake bound (Michaelis–Menten saturation of the FBA upper bound). Table 1 of Harcombe *et al.* [20] lists $K_m = 10$ mM, whereas the Experimental Procedures state $K_m = 0.01$ mM, an internal inconsistency noted in the compiled parameter set. BacArena assigns $K_m$ per exchange reaction through `setKinetics`, using literature transporter values where available (e.g. glucose $K_m = 0.01$ mM in the *P. aeruginosa* biofilm example). PhysiCelldFBA adopts substrate-specific $K_m$ from dynamic-FBA calibrations or reconstruction-consistent assignments (Tables S8, S9, and S10), typically at the 0.001–1 mM scale rather than a single global value.

**Maximum uptake rates ($V_{\max}$).** COMETS sets a uniform $V_{\max} = 10$ mmol $\text{gDW}^{-1}\,\text{h}^{-1}$ for all metabolites. BacArena likewise specifies $v_{\max}$ per exchange reaction (mmol $\text{gDW}^{-1}\,\text{h}^{-1}$) rather than through a global default. PhysiCelldFBA assigns substrate-specific $V^{\text{kin}}_{\max,i}$, scaled to remain compatible with each GEM exchange-reaction upper bound while preserving literature order-of-magnitude where reported; secreted by-products receive $V_{\max}^{\text{kin}} = 0$ so that fermentation products can accumulate in the extracellular field.

### S6. Preprocessing of genome-scale metabolic models

#### S6.1. Model trimming and reduction

This section describes preprocessing steps applied to genome-scale metabolic models before their integration into PhysiCelldFBA, together with the analyses used to select metabolic constraints for specific simulation scenarios.

Before their integration into PhysiCelldFBA, all genome-scale metabolic models were preprocessed to reduce the computational cost of repeatedly solving flux balance analysis (FBA) problems during agent-based simulations while preserving their metabolic phenotype.

Model preprocessing consisted of three steps: (i) harmonisation of exchange reactions, (ii) removal of blocked reactions, and (iii) removal of gap metabolites. Exchange reaction identifiers were first standardised to a common naming convention. Medium constraints were then adjusted to reproduce the reference growth conditions for each organism, and compatible exchange bounds were generated by combining the uptake and secretion capabilities of all models, allowing different organisms to interact through a common extracellular environment.

Blocked reactions were subsequently identified using COBRApy. A reaction (j) is considered blocked if it cannot carry flux in any feasible steady-state solution,

$$v_j = 0 \qquad \forall \mathbf{v} \in \Omega,$$

where

$$\Omega = \{\mathbf{v} \mid S\mathbf{v} = 0,\ \mathbf{l} \le \mathbf{v} \le \mathbf{u}\}$$

denotes the feasible solution space.

Operationally, blocked reactions were detected by independently maximising and minimising the flux through every reaction,

$$\max / \min \quad v_j \text{ subject to} \qquad S\mathbf{v} = 0, \mathbf{l} \le \mathbf{v} \le \mathbf{u}, \tag{15}$$

and classifying reaction (j) as blocked whenever

$$\left|v_j^{\max}\right| < 10^{-7} \quad \text{and} \quad \left|v_j^{\min}\right| < 10^{-7}.$$

Gap metabolites were then identified following the definition of Ponce-de-León et al. [23]. Let

$$R(i) = \{j \mid S_{ij} \neq 0\}$$

be the set of reactions involving metabolite (i). Metabolite (i) is classified as a gap metabolite whenever

$$R(i) \subseteq J_{\text{blocked}},$$

where ($J$_blocked) is the set of blocked reactions. Because neither blocked reactions nor the associated gap metabolites can participate in any feasible metabolic state, they can be removed without altering the model's metabolic capabilities.

To verify that preprocessing preserved the original metabolic phenotype, FBA was performed before and after trimming, and the optimal objective value was required to remain unchanged within a numerical tolerance of ($10^{-7}$). The reduced models were then exported in SBML format and used in all PhysiCelldFBA simulations. By reducing the number of reactions, metabolites, variables, and constraints in the underlying linear programming problem, this preprocessing substantially decreases the computational cost of the metabolic optimisation performed by each cell agent.

#### S6.2. Phenotype phase plane analysis

To select biologically relevant metabolic constraints for the tumour simulations, a phenotype phase plane analysis was performed on the MCF7 metabolic model by systematically varying the maximum oxygen uptake and lactate secretion fluxes while maximising biomass production (Supplementary Figure S7). The resulting phase plane exhibits a broad plateau of near-maximal growth, indicating that similar biomass production rates can be achieved over a wide range of oxygen uptake and lactate secretion fluxes. This extensive plateau reflects the presence of alternative optimal metabolic states within the feasible solution space.

Rather than selecting an arbitrary point within this region, the oxygen uptake constraint used in the spatial simulations was chosen close to the transition between the high-growth plateau and the oxygen-limited regime. At this operating point, relatively small reductions in oxygen availability produce pronounced changes in biomass production and metabolic fluxes. This region is particularly relevant for modelling tumour physiology because oxygen gradients naturally develop around blood vessels, allowing local oxygen depletion to generate distinct proliferative, hypoxic, and necrotic regions as emergent consequences of the metabolic model rather than predefined cellular states.

### Supplementary Note

#### Documentation of the PhysiCelldFBA software package

Comprehensive documentation for PhysiCelldFBA is available online at `https://physicelldfba.github.io/PhysiCelldFBA/`. The documentation includes installation instructions, a step-by-step user guide, detailed descriptions of the PhysiCell–dFBA integration framework, API references, and reproducible example workflows. Tutorial examples cover representative applications of the framework and provide practical guidance for building, configuring, and extending PhysiCelldFBA simulations. The online documentation is actively maintained and contains the most up-to-date information on software features, usage, and examples.

#### Software specification

PhysiCelldFBA is a C++ software extension built on top of PhysiCell v1.14.2. As for other PhysiCell intracellular extensions, it introduces an additional modelling layer while preserving the general structure of the PhysiCell core. In particular, PhysiCelldFBA implements flux balance analysis (FBA) at the single-cell level by defining a new intracellular model class, *dFBAIntracellular*, derived from the PhysiCell *Intracellular* interface. The implementation relies on external numerical and model-parsing libraries: libSBML is used to read SBML models with FBC annotations, whereas the Coin-OR CLP solver is used to formulate and solve the corresponding linear programming problem.

The code specific to PhysiCelldFBA is contained in the directory `addons/dFBA/`. Its main source-code directory, `addons/dFBA/src/`, contains the C++ classes that define the dFBA intracellular model, the metabolic model representation, the reaction and metabolite abstractions, and the data structure used to store FBA solutions. In particular:

**`dfba_intracellular.h` and `dfba_intracellular.cpp`.** These files define and implement the *dFBAIntracellular* class, which is the interface between PhysiCell agents and the intracellular FBA model. The class inherits from PhysiCell's *Intracellular* class and stores the information required to associate a cell with a metabolic model, including the SBML file name, the objective reaction, the current growth rate, the dFBA update time step, and the mapping between PhysiCell diffusive substrates and FBA exchange reactions. The class is also responsible for parsing the dFBA-specific XML configuration blocks.

**`dfba_Model.h` and `dfba_Model.cpp`** These files define the *dFBAModel* class, which represents the intracellular constraint-based metabolic model. The class stores the list of metabolites, the list of reactions, index maps for fast lookup, the current FBA solution, and the Coin-OR CLP linear programming problem. It provides methods to read a metabolic model from an SBML file, extract metabolites and reactions, retrieve FBC flux bounds, assign objective coefficients, and initialise the corresponding stoichiometric linear programming problem.

**`dfba_Reaction.h` and `dfba_Reaction.cpp`** These files define the *dFBAReaction* class. Each object stores the identifier and name of a metabolic reaction, its lower and upper flux bounds, its objective coefficient, its current flux value, and the set of metabolites involved in the reaction together with their stoichiometric coefficients. The class provides methods to determine whether a reaction is reversible, retrieve reactants and products, update bounds and objective coefficients, and return a human-readable reaction string. This class is used by *dFBAModel* to construct the stoichiometric matrix and to store the flux values returned by the solver.

**`dfba_Metabolite.h` and `dfba_Metabolite.cpp`** These files define the *dFBAMetabolite* class. This is a lightweight data structure used to store the identifier and name of each metabolite read from the SBML model. The metabolite objects are indexed by *dFBAModel* and used when constructing reactions and the stoichiometric matrix.

**`dfba_Solution.h` and `dfba_Solution.cpp`** These files define the *dFBASolution* class, which stores the output of an FBA optimisation. The solution object contains the objective value, the solver status, the flux distribution indexed by reaction identifier, and reduced costs. It provides a compact container through which the metabolic state of a cell can be accessed after each FBA update.

#### Installation guide

PhysiCelldFBA is distributed through GitHub and can be installed in the same way as a standard PhysiCell distribution. The repository can be cloned with:

```
git clone https://github.com/PhysiCelldFBA/PhysiCelldFBA.git
```

After cloning the repository, the user can compile one of the dFBA sample projects contained in:

```
sample\_projects\_intracellular/fba/
```

These projects follow the same general organisation as standard PhysiCell sample projects, but include a custom `Makefile` that adds the dFBA source files, header paths, and external libraries required by PhysiCelldFBA. The dFBA `Makefile` includes an automatic first-time setup script for the external libraries. For a selected dFBA sample project, the target `libFBA` points to the expected Coin-OR header file:

```
/coin-or/include/coin/CoinPackedMatrix.hpp
```

If this file is not present, the `Makefile` runs:

```
python3 beta/setup_fba.py
```

The script `beta/setup_fba.py` determines the operating system and architecture, creates the external-library directory if needed, and installs the required `coin-or` and `libsbml` packages under `addons/dFBA/ext`. The package URLs are specified in `beta/fba_packages.json`. The setup script checks whether each package directory already exists and is populated; if so, the package is considered already installed and the download is skipped. Therefore, this setup step is normally triggered only during the first compilation of a dFBA project, or whenever the external-library directory has been removed.

In addition to the standard PhysiCell input files, such as `PhysiCell_settings.xml`, `cells.csv`, and other project-specific configuration files, PhysiCelldFBA requires at least one SBML model file. The SBML model is associated with a dFBA cell definition through the intracellular configuration block and provides the metabolic network on which flux balance analysis is performed.

#### dFBA cell definition block

To assign a dFBA model to a PhysiCell cell definition, the user must add an `intracellular` block with `type="dfba"` inside the corresponding `cell_definition` in `PhysiCell_settings.xml` (See Figure S2). This block specifies the SBML metabolic model associated with the cell, the time step used to update the intracellular FBA model, the mapping between extracellular substrates and FBA exchange reactions, and the parameters used to convert the FBA solution into cell growth and phenotype updates.

The `settings` block defines the SBML file used by the cell and the frequency at which the intracellular FBA model is updated. The `sbml_filename` entry must point to a valid SBML-FBC model file, usually located in the project `config/` directory. The `intracellular_dt` entry sets the dFBA update interval in minutes. In order to avoid artifacts in the simulation, it is optimal to match the intracellular_dt with the diffusion_dt.

The `transport_model` block defines how extracellular substrates in the PhysiCell microenvironment are connected to exchange reactions in the FBA model. Each `exchange` entry links one PhysiCell substrate, specified by the `substrate` attribute, to one FBA exchange reaction, specified by `fba_flux`. The substrate name must match a substrate defined in the PhysiCell microenvironment, and the FBA flux identifier must match a reaction identifier in the SBML model. The parameters `Km` and `Vmax` define the kinetic relation used to constrain the corresponding exchange flux as a function of the local extracellular substrate concentration. Multiple `exchange` entries can be included, one for each extracellular substrate coupled to the metabolic model.

The `growth_model` block defines how the optimised metabolic objective is interpreted as cell growth. The parameter `cell_density` is used to convert between cell volume and biomass, `reference_volume` defines the reference cell volume used for growth and division, and `max_growth_rate` sets an upper bound for the growth rate imposed by the intracellular model. The `objective_reaction` entry specifies the reaction whose flux is interpreted as the metabolic objective for growth, and must correspond to a valid reaction identifier in the SBML model.

The optional `death_model` block defines whether metabolic-dependent cell death is enabled. If enabled, `death_type` specifies the PhysiCell death process to activate (apoptosis or necrosis), `death_trigger_flux` identifies the FBA reaction used as the death criterion, `death_flux_threshold` defines the threshold value for that flux, and `death_rate_increase` defines the increase in the corresponding death rate.

#### Auxiliary scripts for PhysiCelldFBA configuration file generation

##### Overview of the `generate_dfba_yaml.py` workflow

The `generate_dfba_yaml.py` script implements an automated pipeline for generating PhysiCelldFBA configuration files. This utility addresses a critical bottleneck in hybrid simulation setup: the manual specification of mappings between extracellular substrates tracked by the physics-based agent model of PhysiCell and intracellular fluxes computed by the dynamic Flux Balance Analysis (dFBA) solver. Here we will detail the functioning of this script.

**a. Input Requirements and Validation** The script accepts three primary inputs:

1. **SBML-encoded metabolic model** (`--sbml`): A constraint-based reconstruction in standard SBML format, typically a GEM such as Recon3D. The parser validates model completeness by confirming the presence of (i) at least one reaction annotated as biomass synthesis, (ii) exchange reactions following the conventional `EX_` prefix nomenclature, and (iii) defined flux bounds for all reactions.
2. **Substrate mapping file** (`--substrates`): A comma-separated value file establishing bidirectional correspondence between PhysiCell microenvironment variable names (e.g., `oxygen`, `glucose`) and their cognate metabolite identifiers in the GEM (e.g., `M_o2_e`, `M_glc__D_e`). This mapping is essential because PhysiCell tracks substrates using descriptive names in its diffusion solver, whereas FBA models employ metabolite identifiers derived from biochemical databases (BiGG, MetaNetX).
3. **Biomass reaction identifier** (`--biomass_rxn`): Explicit specification of the biomass pseudoreaction, as automated detection remains unreliable across heterogeneous model repositories due to inconsistent annotation practices.

**b. Core Processing Logic** **b.1. Exchange Reaction Extraction and Classification:** The script parses the SBML model using the `libsbml` Python bindings and identifies all exchange reactions through pattern matching on reaction identifiers. Each exchange reaction is classified as either:

- **Import** (substrate uptake): Flux bounds permit negative values, indicating net influx into the intracellular compartment.
- **Export** (byproduct secretion): Flux bounds permit positive values, indicating net efflux.

This classification determines the directionality of substrate coupling in the subsequent YAML output.

**b.2. Flux Bound Harmonisation:** A critical preprocessing step involves reconciling units between the metabolic model and the agent-based framework. PhysiCelldFBA internally normalises fluxes to mmol/gDW/hr, but GEMs frequently employ alternative units (e.g., mmol/gDCW/hr, $\mu$mol/$10^6$ cells/hr). The script applies configurable scaling factors (`--flux_scaling`) to transform upper and lower bounds into the expected unit system. Explicit warnings are emitted when bounds exceed physiologically plausible ranges (default threshold: $|\text{bound}| > 1000$ mmol/gDW/hr).

**b.3. Substrate-Metabolite Linkage Construction:** For each mapped substrate, the script:

1. Locates the corresponding exchange reaction via the metabolite identifier.
2. Extracts the default bounds and objective coefficient (typically zero for exchange reactions).
3. Stores the PhysiCell substrate name as the coupling key, enabling the runtime solver to query substrate concentrations from the microenvironment and dynamically adjust exchange reaction bounds accordingly.

**b.4. YAML Structure Generation:** The output YAML adheres to a hierarchical schema of the XML (Figure S2) with the following top-level keys:

The `default_bound` parameter preserves the original model bound as a fallback when substrate concentrations fall below the coupling threshold, preventing numerical instabilities at vanishingly low concentrations.

**c. Integration with the PhysiCelldFBA Runtime.** At simulation initialisation, the PhysiCelldFBA core reads the generated YAML and constructs an internal mapping data structure enabling $O(1)$ lookup of exchange reactions by substrate name. During each agent's dFBA solve (executed at user-specified intervals via the `dFBA_dt` parameter), the solver:

1. Queries the local substrate concentration, $[S]$, from the PhysiCell microenvironment.

2. Applies a Michaelis-Menten-style scaling function to compute effective bounds:

$$v_{\max}^{\text{eff}} = v_{\max} \cdot \frac{[S]}{K_m + [S]} \tag{16}$$

3. Substitutes the effective bounds into the linear programming (LP) constraint matrix.

4. Solves for the optimal flux distribution maximising the biomass objective.

**d. Command-Line Interface** A typical execution requires specifying the SBML model, the substrate map, and the biomass reaction:

```
python generate_dfba_yaml.py \
  --sbml model/recon3d.xml \
  --substrates config/substrate_map.csv \
  --biomass_rxn BIOMASS_human \
  --output physicell_dfba_config.xml \
  --flux_scaling 1.0 \
  --solver glpk
```

Full option documentation is accessible via the `--help` flag. Example configuration files for several standard GEMs are provided in the repository's `examples/` directory.

##### Role of the `dfba_configurator.py` Utility

While the `generate_dfba_yaml.py` script establishes the foundational metabolic constraints and substrate mappings, the `dfba_configurator.py` script acts as the critical integration layer between these metabolic definitions and the overarching PhysiCell agent-based model. Because PhysiCell relies on a specific XML schema to define cellular phenotypes and the diffusive microenvironment, the dFBA configuration cannot exist as an isolated file; it must be embedded within the master PhysiCell configuration file (typically `PhysiCell_settings.xml`). The configurator automates this injection process, preventing manual XML editing errors and ensuring that the C++ runtime initialises the dFBA solvers with the correct pointers to both the metabolic models and the microenvironmental substrate fields.

**Microenvironment Schema Synchronisation** A prerequisite for PhysiCelldFBA simulations is that every extracellular substrate tracked by the BioFVM solver must have a corresponding diffusive field defined in the PhysiCell microenvironment block. The `dfba_configurator.py` script parses the generated dFBA YAML to extract all mapped substrate names and cross-references them against the `<microenvironment_variables>` in the input XML. If a substrate required by the metabolic model (e.g., `glucose`) is absent from the XML, the script dynamically injects it. During this injection, it assigns default biophysical properties (specifically the diffusion coefficient ($D$) and the initial uniform concentration) either from a user-supplied dictionary or from physiologically informed defaults. This automated schema synchronisation prevents the segmentation faults and null-pointer exceptions that typically occur when the C++ backend attempts to query a non-existent substrate field during the flux balance calculation.

**Cell-Definition Integration and Multi-Lineage Assignment** To support the modelling of heterogeneous tissues, PhysiCell structures agents via `<cell_definition>` blocks, which dictate phenotype-specific behaviours such as cycle progression and motility. The configurator script identifies target cell definitions by user-specified names (e.g., `tumour_cell`, `macrophage`) and injects a nested `<dFBA>` XML node into each. Within this node, the script translates the hierarchical key-value pairs of the YAML file into XML attributes. Crucially, this allows for multi-lineage metabolic modelling: the configurator can assign entirely distinct GEMs to different cell types by pointing separate cell definitions to different YAML outputs, or it can assign the same GEM with cell-type-specific flux bounds, reflecting known physiological differences in uptake rates between cell types.

**Parameter Hierarchy and Local Overrides** The configurator implements a hierarchical logic for parameter resolution. Global parameters defined at the root of the dFBA YAML such as the linear programming solver tolerance, the maximum iteration count, or the global coupling half-saturation constant ($K_m$), are applied as baseline defaults to all cell definitions. However, the script allows for local overrides within the XML injection step. If a specific cell type requires a distinct kinetic coupling to a substrate (e.g., a highly glycolytic cell line with an elevated $v_{\max}$ for glucose uptake), the configurator writes these local parameters directly into the cell-specific `<dFBA>` block. At runtime, the C++ dFBA engine evaluates this hierarchy, prioritising cell-level bounds over global YAML bounds, thereby enabling the specification of intercellular metabolic heterogeneity without the need to maintain duplicate, redundant metabolic models.

**Execution and Downstream Compatibility** The script is executed as a command-line utility that takes the path to the base PhysiCell XML, the path to the dFBA YAML, and an output path for the unified configuration file. By abstracting the XML manipulation into this Python-based preprocessing step, the PhysiCelldFBA framework maintains backward compatibility with standard PhysiCell parsers and the PhysiCell Studio GUI, which we discuss in the following section. Users can continue to use visual tools to adjust mechanical or cycle parameters in the XML; upon re-running the configurator, the dFBA nodes are seamlessly updated or re-injected without corrupting the surrounding agent-based configuration data. A typical execution to couple a previously generated YAML to a base PhysiCell configuration is structured as follows:

```
python dfba_configurator.py \
  --config PhysiCell_settings.xml \
  --dfba_yaml my_model.yaml \
  --output PhysiCell_settings_with_dFBA.xml \
  --cell_types tumour_cell,macrophage
```

Here, the `--cell_types` flag explicitly dictates which cellular phenotypes receive the `<dFBA>` node injection, ensuring that non-metabolically active agent types remain unburdened by solver overhead.

#### PhysiCell-Studio compatibility

PhysiCelldFBA is compatible with PhysiCell-Studio, the graphical user interface developed for PhysiCell projects. Users can clone PhysiCell-Studio from:

```
git clone https://github.com/PhysiCell-Tools/PhysiCell-Studio.git
```

and use it to open, edit, run, and save PhysiCelldFBA projects in the same way as standard PhysiCell or PhysiBoSS projects (Figure S4).

PhysiCell-Studio includes support for the dFBA intracellular model through the `Intracellular` tab. A user does not need to manually write the dFBA block in `PhysiCell_settings.xml` before opening the project. Instead, the user can select `dFBA` from the list of available intracellular model types and fill in the corresponding fields in the graphical interface. These fields define the SBML file, the dFBA update time step, the transport model, the growth model, and the optional metabolic-dependent death model.

When the project is saved or executed from PhysiCell-Studio, the graphical interface writes the corresponding `<intracellular type="dfba">` block into the configuration file using the correct XML structure. This allows users to generate a valid dFBA cell-definition block without manually editing the XML file.

The interface also helps reduce configuration errors. In particular, the transport model is edited through a table in which each exchange reaction is associated with a PhysiCell substrate. The substrate field is selected from the list of substrates defined in the microenvironment, which helps enforce consistency between the substrate names used by PhysiCell and the exchange fluxes specified for the FBA model. The remaining fields allow the user to specify the FBA exchange reaction identifier, the kinetic parameters `Km` and `Vmax`, and the growth and death-model parameters required by PhysiCelldFBA.

Thus, PhysiCell-Studio provides a graphical workflow for creating and editing PhysiCelldFBA models while preserving compatibility with the underlying `PhysiCell_settings.xml` format.

### Supplementary Tables

Table S12: Symbols used throughout Supplementary Methods.

| Symbol | Meaning | Units |
|---|---|---|
| $[S_i]$ | Extracellular concentration of substrate $i$ | mM |
| $D_i$ | Diffusion coefficient of substrate $i$ | $\mu$m$^2$ min$^{-1}$ |
| $\lambda_i$ | First-order decay rate | min$^{-1}$ |
| $\mathbf{x}$ | Spatial position | $\mu$m |
| $\mathbf{x}_k$ | Position of cell $k$ | $\mu$m |
| $V_{voxel}$ | BioFVM voxel volume | $\mu$m$^3$ |
| $\alpha$ | Conversion from $\mu$m$^3$ to litres | $10^{-15}$ L $\mu$m$^{-3}$ |
| $M_{voxel,i}$ | Mass of substrate $i$ contained in one voxel | mmol |
| $V_{cell}$ | Total cell volume | $\mu$m$^3$ |
| $V_{solid}$ | Cell solid volume | $\mu$m$^3$ |
| $\phi_{fluid}$ | Cellular fluid fraction | dimensionless |
| $\rho_{cell}$ | Cellular dry-mass density | pg $\mu$m$^{-3}$ |
| $m_{DW}$ | Cell dry weight | g |
| $\mathbf{N}$ | Stoichiometric matrix | – |
| $\mathbf{v}$ | Metabolic flux vector | mmol gDW$^{-1}$ h$^{-1}$ |
| $\mathbf{c}$ | Objective coefficient vector | – |
| $Z$ | FBA objective value | – |
| $\mu$ | Specific growth rate | h$^{-1}$ |
| $v_i$ | Exchange flux for substrate $i$ | mmol gDW$^{-1}$ h$^{-1}$ |
| $v_{max,i}$ | Michaelis–Menten uptake limit | mmol gDW$^{-1}$ h$^{-1}$ |
| $V_{max,i}^{kin}$ | Maximum transport rate parameter | mmol gDW$^{-1}$ h$^{-1}$ |
| $K_{m,i}$ | Michaelis constant | mM |
| $v_{limit,i}$ | Maximum physically feasible uptake | mmol gDW$^{-1}$ h$^{-1}$ |
| $v_{lb,i}$ | Exchange lower bound applied to FBA | mmol gDW$^{-1}$ h$^{-1}$ |
| $E_i^k$ | Net export rate of substrate $i$ by cell $k$ | mmol min$^{-1}$ |
| $\Delta t_{diff}$ | Diffusion timestep | min |
| $\Delta t_{intracellular}$ | Intracellular/FBA timestep | min |

#### Supplementary Figures

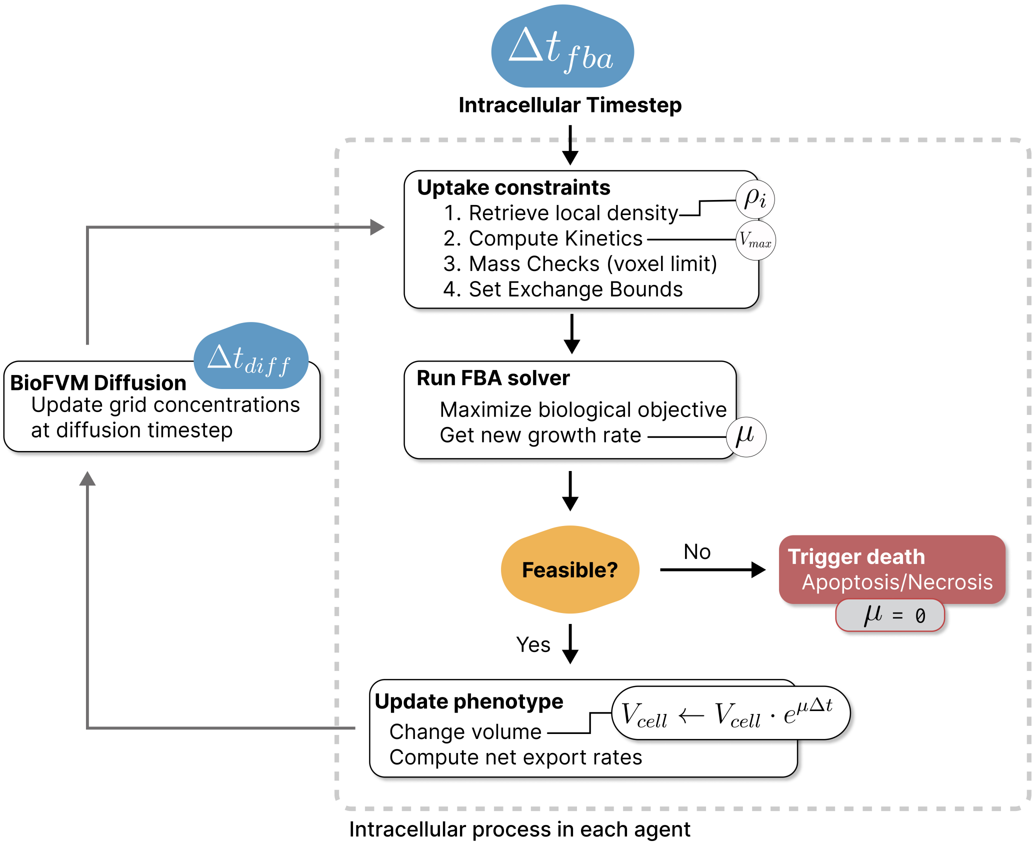

Figure S1: Overview of the PhysiCelldFBA workflow executed at each intracellular timestep.

```
<intracellular type="dfba">
    <settings>
        <sbml_filename>./config/Ecoli_core.xml</sbml_filename>
        <intracellular_dt>0.01</intracellular_dt>
    </settings>
    <transport_model>
        <exchange substrate="glucose">
            <fba_flux>R_EX_glc__D_e</fba_flux>
            <Km units="mM">0.02</Km>
            <Vmax units="mmol/gDW cell/hour">8.0</Vmax>
        </exchange>
        <exchange substrate="acetate">
            <fba_flux>R_EX_ac_e</fba_flux>
            <Km units="mM">1</Km>
            <Vmax units="mmol/gDW cell/hour">0.0</Vmax>
        </exchange>
        <exchange substrate="oxygen">
            <fba_flux>R_EX_o2_e</fba_flux>
            <Km units="mM">0.002</Km>
            <Vmax units="mmol/gDW cell/hour">18</Vmax>
        </exchange>
        ...
    </transport_model>
    <growth_model>
        <objective_reaction>R_BIOMASS_Ecoli_core_w_GAM</objective_reaction>
        <max_growth_rate units="1/hours">0.8</max_growth_rate>
        <reference_volume units="micron^3">1.3</reference_volume>
        <cell_density units="g/ml">1.04</cell_density>
    </growth_model>
</intracellular>
```

Figure S2: Example of an intracellular dFBA configuration for an *E. coli* cell definition.

```yaml
substrates:
  - name: "glucose"
    diffusion_coefficient: 50000.0  # micron^2/min
    decay_rate: 0.0                  # 1/min
    initial_condition: 0.0            # mM (optional, defaults to 0.0)

  - name: "acetate"
    diffusion_coefficient: 50000.0
    decay_rate: 0.0
    initial_condition: 0.0

  - name: "oxygen"
    diffusion_coefficient: 100000.0
    decay_rate: 0.1
    initial_condition: 0.0

  - name: "CO2"
    diffusion_coefficient: 100000.0
    decay_rate: 0.0
    initial_condition: 0.0

models:
  # Model 1: iML1515
  iML1515:
    # Path to SBML file (can be relative to --sbml-folder or absolute)
    sbml_path: "iML1515.xml"

    exchanges:
      - substrate: "glucose"           # Substrate name (must match microenvironment variable)
        fba_flux: "R_EX_glc__D_e"      # Exchange reaction ID from SBML model
        Km: 0.01                       # Michaelis-Menten constant (mM)
        Vmax: 10.0                     # Maximum velocity (mmol/gDW cell/hour)

      - substrate: "acetate"
        fba_flux: "R_EX_ac_e"
        Km: 0.01
        Vmax: 10.0

      - substrate: "oxygen"
        fba_flux: "R_EX_o2_e"
        Km: 0.01
        Vmax: 10.0

      - substrate: "CO2"
        fba_flux: "R_EX_co2_e"
        Km: 0.01
        Vmax: 10.0

    growth_model:
      cell_density: 1.04              # Cell density (g/ml)
      reference_volume: 1.3           # Reference volume for cell division (um^3)
      nuclear_volume: 0.0             # Nuclear volume (um^3)
      max_growth_rate: 0.95           # Maximum growth rate (1/h)
      objective_reaction: "R_BIOMASS_Ec_iML1515_core_75p37M"  # Objective reaction ID from SBML model

    settings:
      intracellular_dt: 0.01          # Intracellular time step (min)

    death_model:
      enabled: true
      death_type: necrosis           # death type string
      death_trigger_flux: "R_ATPM"  # mmol/gDW cell/hour
      death_flux_threshold: 8.4     # dimensionless
      death_rate_increase: 1.67e-5  # rate increase
```

Figure S3: YAML dFBA configuration for a given cell definition.

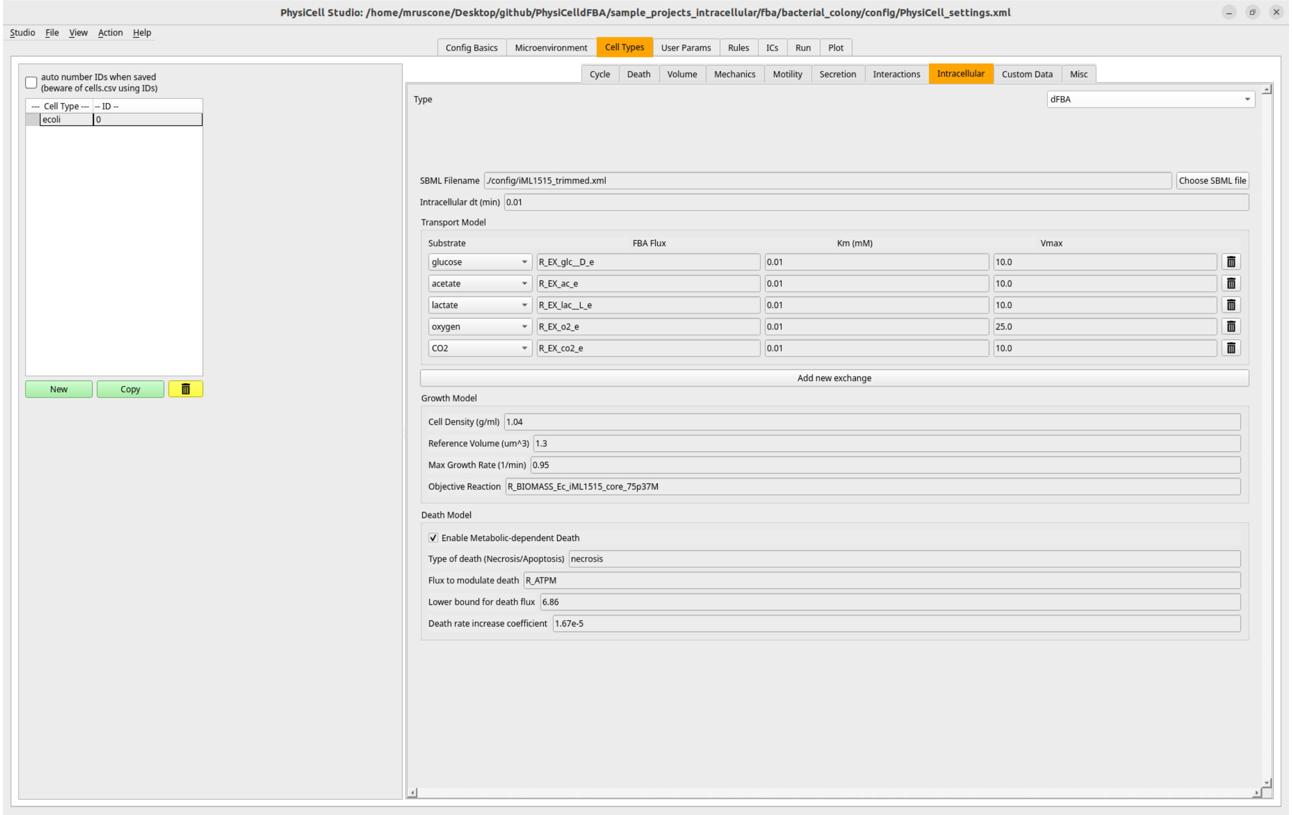

Figure S4: PhysiCell-Studio interface for configuring the dFBA intracellular model. The `Intracellular` tab allows the user to select `dFBA`, choose the SBML model, define the intracellular update time step, map PhysiCell substrates to FBA exchange reactions, and specify growth and optional metabolic-dependent death parameters.

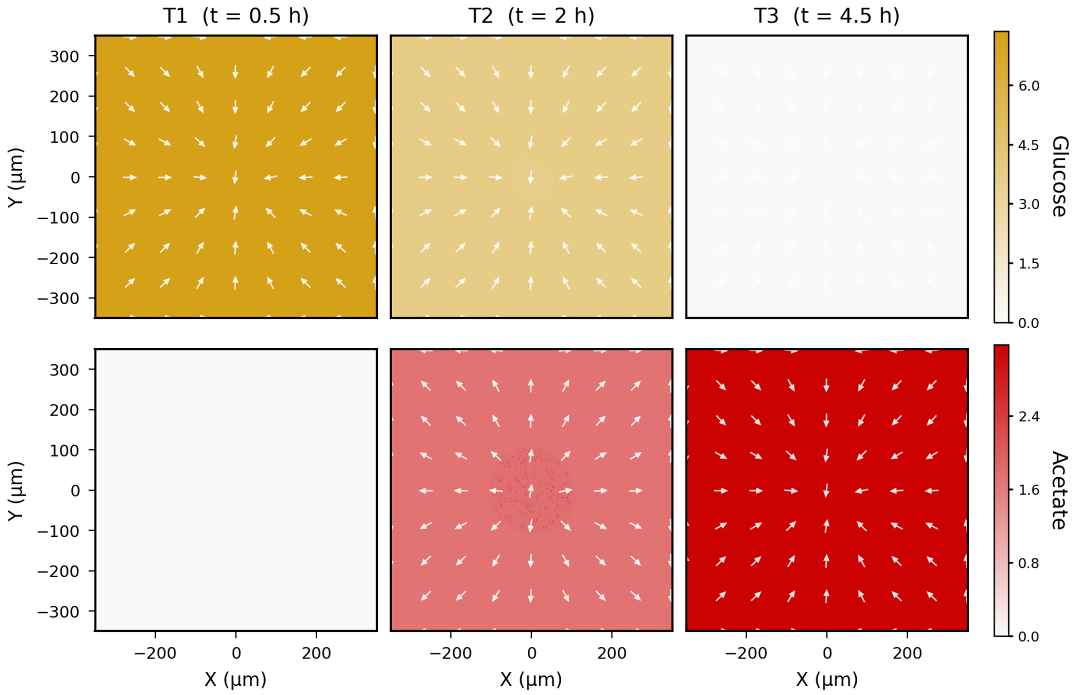

Figure S5: *Escherichia coli* acetate switch: sequential substrate consumption from overflow metabolism. Spatiotemporal snapshots of *E. coli* growth on a mixed-substrate environment over 10 hours of simulated time, showing the diauxic shift from glucose utilisation to acetate consumption. The simulation demonstrates overflow acetate secretion during oxygen-limited growth on glucose and the subsequent re-consumption of accumulated acetate after glucose depletion, a classic example of hierarchical substrate utilisation in *E. coli*. This scenario validates the PhysiCelldFBA coupling by showing that the dynamic flux balance analysis correctly captures the metabolic transition between glucose and acetate oxidation, with the timing and magnitude of the switch determined by intracellular metabolic constraints rather than explicit phenotypic instructions.

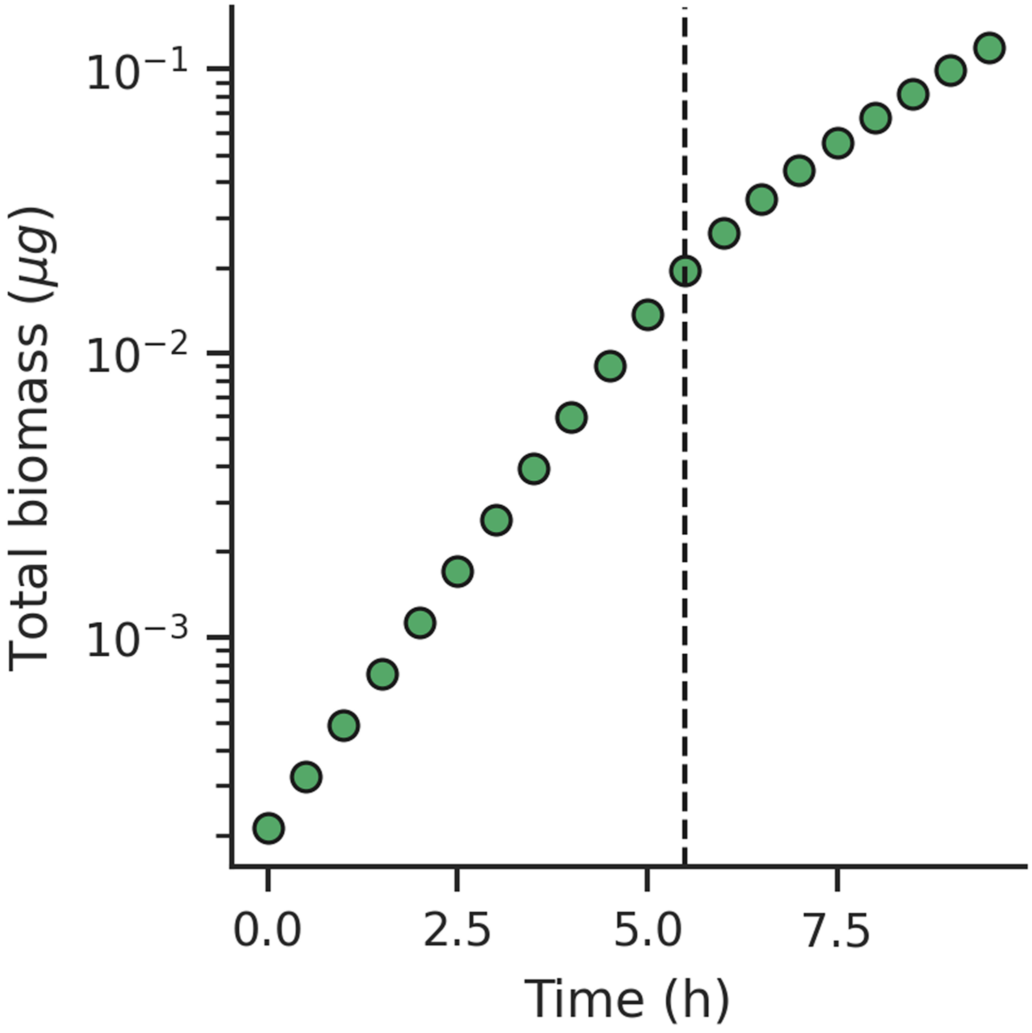

Figure S6: Quantification of the transition from exponential to sub-exponential growth in the simulated *E. coli* colony. Total colony biomass (log scale) as a function of simulated time shows near-exponential growth during the early phase of colony expansion, followed by a marked deceleration as nutrient availability becomes limiting in the colony interior. The dashed vertical line marks $t^* \approx 5.5$ h, the transition time estimated by fitting two independent linear regressions to ln(biomass) on either side of each candidate breakpoint and selecting the breakpoint that minimises the combined residual sum of squares.

To pinpoint when colony growth shifted from exponential to sub-exponential, we fit two straight lines to the log-transformed biomass curve: one for an early period, one for a later period, and tested every possible split point between them. The split point where the two lines together fit the data best was taken as the transition time, $t^* \approx 5.5$ h. Before this point, biomass doubled roughly every 0.84 h; afterwards, doubling slowed to about 1.56 h. As an independent check, we also tracked the growth rate directly at each time point (rather than assuming two distinct phases); it stayed roughly flat early on and then declined smoothly around $t^*$, matching the fitted transition.

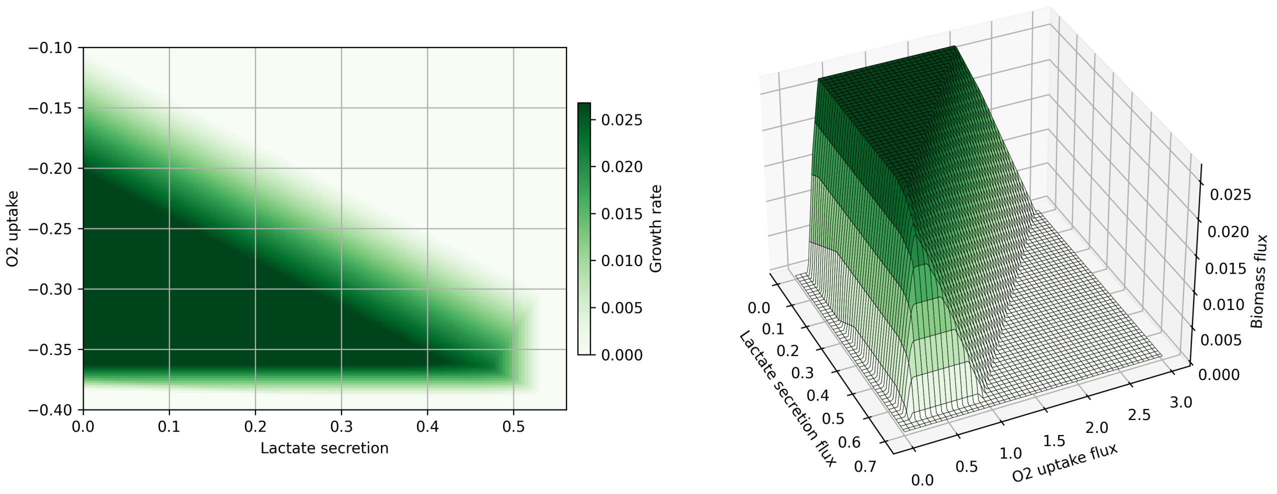

Figure S7: Phenotypic phase plane analysis of the MCF7 metabolic model: biomass production as a function of maximum oxygen uptake and lactate secretion constraints. (Left) Two-dimensional heatmap showing growth rate (colour intensity, green scale: 0.000–0.025 $h^{-1}$) across the metabolic flux space defined by oxygen uptake (y-axis, mmol·$min^{-1}$, shown as negative indicating consumption) and lactate secretion (x-axis, mmol·$min^{-1}$). (Right) Three-dimensional surface representation of the same phenotypic landscape. The phase plane reveals a broad plateau of near-maximal biomass production across a wide range of oxygen and lactate flux combinations, reflecting the presence of multiple alternative optimal metabolic states within the feasible solution space. This plateau is particularly relevant for spatial tumour simulations: the oxygen uptake constraint used in the tissue model (approximately at the transition between the high-growth plateau and the oxygen-limited regime) ensures that small reductions in local oxygen availability produce pronounced changes in biomass production and metabolic fluxes, allowing oxygen gradients to naturally generate distinct proliferative, hypoxic, and necrotic zones as emergent consequences of metabolic constraints rather than predefined phenotypic states.

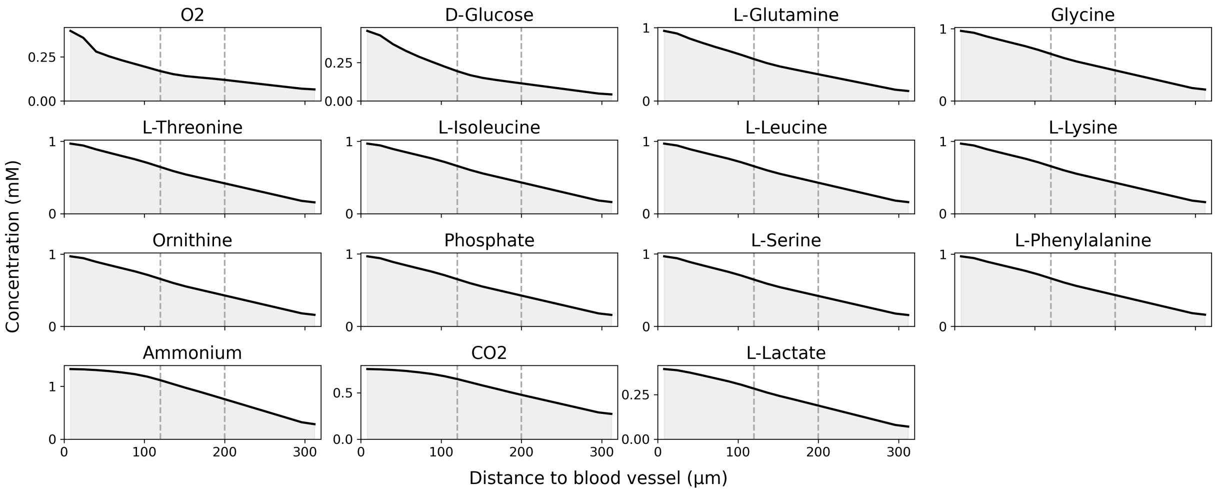

Figure S8: Spatially-resolved metabolite concentration gradients in simulated cancer tissue. Extracellular metabolite concentration profiles are shown as a function of distance from the blood vessel source (at $x = 0$) at 72 hours of simulation time. The grid displays concentration profiles for 15 exchanged metabolites: oxygen ($O_2$), glucose (D-Glucose), glutamine (L-Glutamine), glycine, threonine, isoleucine, leucine, lysine, ornithine, phosphate, serine, phenylalanine, ammonium, carbon dioxide ($CO_2$), and lactate (L-Lactate). Spatial substrate gradients arise from the competing processes of diffusion from the nutrient-rich vasculature and cell-level consumption, resulting in progressive depletion moving away from the vessel. These concentration profiles constrain the metabolic phenotypes of cells in different tissue regions, mechanistically linking molecular transport to intracellular FBA and cell fate decisions.

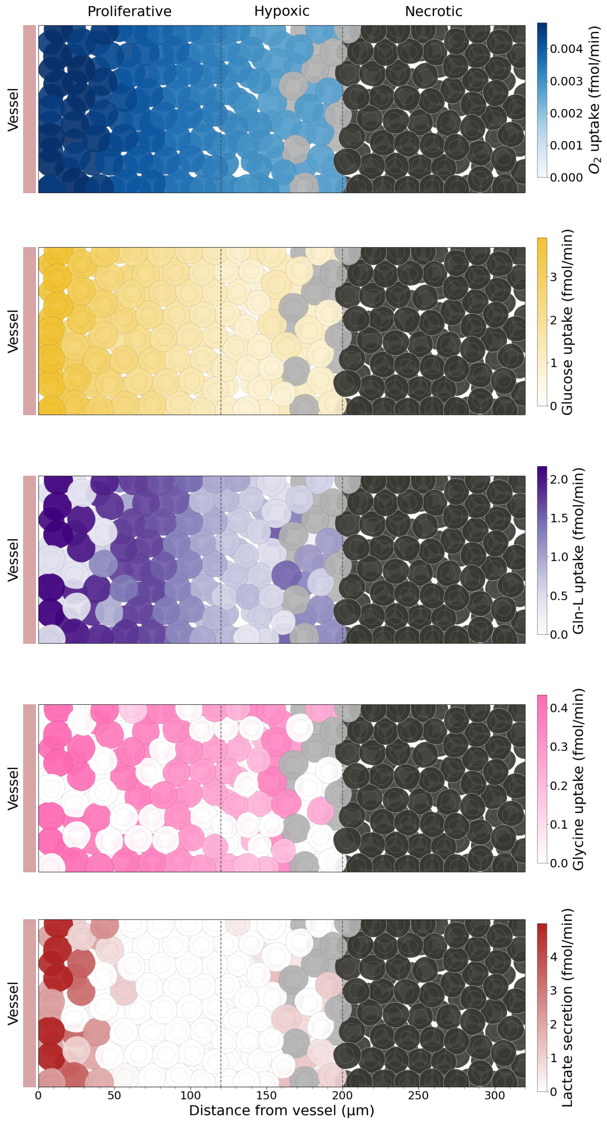

Figure S9: Metabolic heterogeneity and phenotype-specific nutrient uptake at 72 hours. Spatial heatmaps show substrate-specific uptake rates across the simulated cancer tissue at 72 hours, stratified by phenotypic zone. Five key metabolic processes are visualised: oxygen ($O_2$) uptake (blue), glucose uptake (yellow), glutamine uptake (purple), glycine uptake (pink), and lactate secretion (red). Individual cells are rendered as circles with colour intensity indicating uptake flux magnitude in fmol$\cdot$min$^{-1}$. The tissue exhibits three distinct phenotypic regions: proliferative cells adjacent to the vessel (left, high nutrient influx), hypoxic cells in the intermediate zone (centre, reduced oxygen and glucose availability), and necrotic cells in the nutrient-depleted interior (right, minimal metabolic activity). This spatial organisation demonstrates how intracellular metabolic solutions computed via FBA automatically integrate local constraints imposed by spatial substrate availability, creating emergent zones of metabolic specialisation without explicit zone-specific instructions.

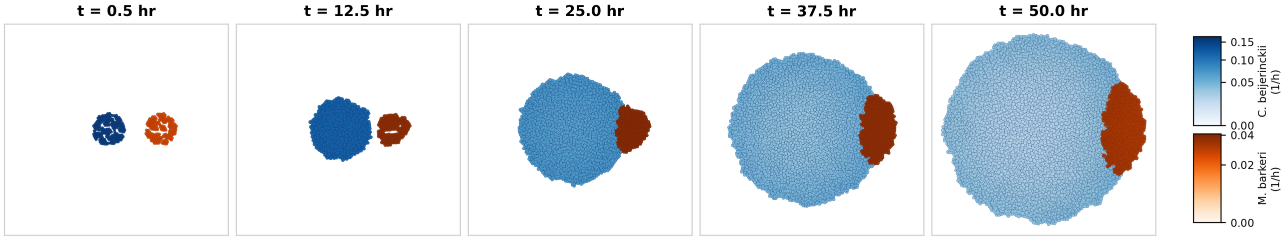

Figure S10: Compact time series of spatial growth-rate dynamics in the syntrophic consortium. Each panel shows the shared microenvironment at a fixed time point (0.5, 12.5, 25.0, 37.5, and 50.0 h). Individual cells are coloured by instantaneous growth rate ($h^{-1}$): *Clostridium beijerinckii* (CB) in blue and *Methanosarcina barkeri* (MB) in orange, using the species-specific colour scales indicated at right (CB: 0.00–0.15 $h^{-1}$; MB: 0.00–0.04 $h^{-1}$). At early times, the two populations remain spatially separated while both sustain positive growth. As the simulation progresses, CB expands more rapidly under glucose supply, the colonies make contact, and MB becomes localised at the CB interface. By 50 h, CB dominates the biomass and exhibits a growth-rate gradient with higher rates at the colony periphery and lower rates in the interior, while MB persists as a spatially confined subpopulation at the colony edge. These snapshots complement Figure 5-E by showing the full temporal progression of spatial niche formation and growth-rate heterogeneity driven by metabolic cross-feeding.

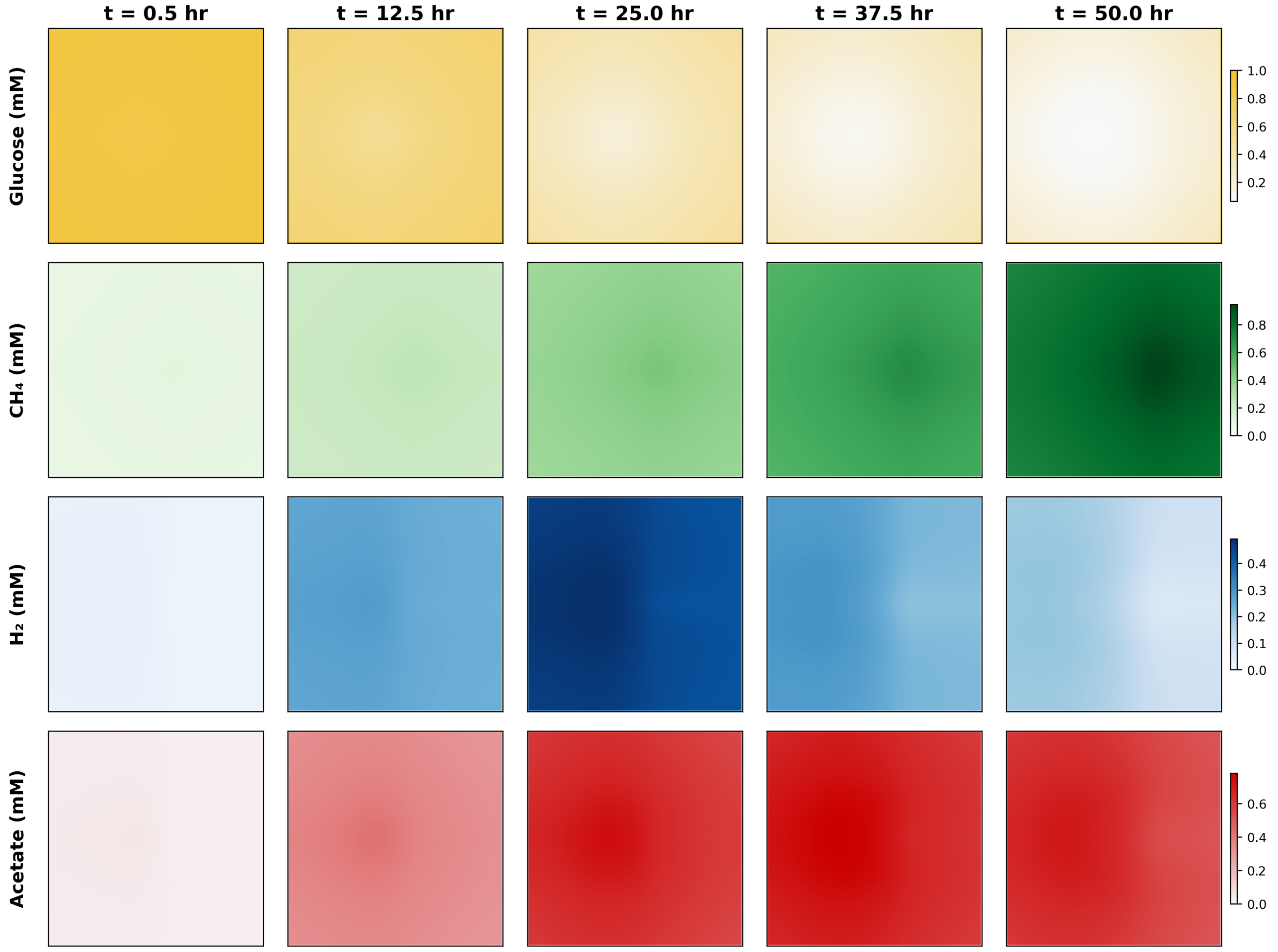

Figure S11: Compact time series of spatial substrate-field dynamics in the syntrophic consortium. Each column shows the shared microenvironment at a fixed time point (0.5, 12.5, 25.0, 37.5, and 50.0 h); each row shows the spatial concentration field of a key metabolite (mM): glucose (top), methane ($CH_4$), hydrogen ($H_2$), and acetate (bottom), using the substrate-specific colour scales indicated at right. At early times, glucose is uniformly available and both populations grow in spatially separated niches. As CB expands and consumes glucose, a depletion zone develops in the colony interior, while $H_2$ and acetate accumulate in the CB-dominated region as fermentative by-products. MB subsequently consumes these cross-fed metabolites, producing a localised $CH_4$-rich zone at the colony interface. By 50 h, glucose is largely depleted in the central domain, acetate and transient $H_2$ gradients reflect diffusion-limited exchange between producer and consumer populations, and methane production is confined to the MB-associated subpopulation. These snapshots complement Figure 5-E by showing the full temporal progression of substrate microgradient formation driven by metabolic coupling, rather than initial and final states alone.

### Supplementary Videos

**Supplementary Video S1.** Temporal evolution of acetate exchange fluxes in an expanding *E. coli* colony. Animated visualisation of single-cell acetate exchange fluxes during colony growth, with cells positioned according to their simulated spatial coordinates and colour indicating the instantaneous acetate exchange rate. As the colony expands and nutrient gradients develop, acetate metabolism becomes spatially segregated: peripheral cells predominantly secrete acetate as a by-product of overflow metabolism, whereas cells in the nutrient-limited interior progressively reduce secretion and re-assimilate acetate. The resulting spatial pattern reveals a division of metabolic labour, with acetate produced by peripheral cells becoming available as a carbon source for cells deeper within the colony. This cross-feeding structure emerges from local differences in substrate availability and metabolic optimisation, without imposing distinct metabolic phenotypes or spatial organisation a priori.

**Supplementary Video S2.** Temporal evolution of cell growth rates in simulated 3D cancer tissue. Animated visualisation of growth rate heterogeneity over 72 hours of simulated tissue expansion. Cell-level growth rates, computed as the FBA-predicted biomass production flux scaled to cell volume changes per unit time, are colour-coded and rendered as individual cell circles arranged in their simulated spatial positions. The animation reveals the progressive establishment of phenotypic zonation: rapid growth of cells near the nutrient source (warm colours, high growth rate) contrasts sharply with growth arrest and cell death in nutrient-depleted interior regions (cool/dark colours). This temporal dynamic demonstrates how the multiscale coupling between extracellular diffusion, intracellular metabolic optimisation, and cell phenotype creates self-organised spatial structure. The pattern emerges purely from local constraints on substrate availability and the cell's adaptive response via metabolic reprogramming.

**Supplementary Video S3.** Temporal evolution of metabolite-specific uptake rates across phenotypic zones. Animated time course of substrate-specific uptake rates across the simulated tissue over 72 hours. The visualisation shows five metabolic processes—oxygen, glucose, glutamine, and glycine uptake, plus lactate secretion—colour-coded across the proliferative, hypoxic, and necrotic zones. Early in the simulation, cells uniformly access abundant nutrients near the vessel. As tissue expands and substrate gradients develop, distinct metabolic zones emerge: proliferative cells near the vessel exhibit high uptake of aerobic substrates; hypoxic cells display reduced but sustained glucose and glutamine uptake; necrotic cells show minimal metabolic activity. This animation illustrates the feedback between tissue growth dynamics, local nutrient depletion, and metabolic state transitions, demonstrating how spatial metabolic heterogeneity emerges endogenously from the spatiotemporal dynamics of transport and consumption.

**Supplementary Video S4.** Time-lapse visualisation of the syntrophic consortium (composed by *Clostridium beijerinckii* and *Methanosarcina barkeri*) over the 50 h simulation. Each frame shows the two-species spatial layout in the shared microenvironment; cell position is displayed in $\mu$m and cell colour encodes species-specific instantaneous growth rate ($h^{-1}$), using the colour scales shown in the figure legend (blue, CB; orange, MB). On-screen counters report live population sizes for CB, MB, apoptotic, and necrotic cells at each time point. The movie captures the faster expansion of CB under glucose supply, the slower but sustained growth of MB at the colony interface, and the emergence of spatial growth-rate gradients as cross-fed metabolites ($H_2$, acetate) are produced, diffuse, and consumed. This animation complements Figure 5-E by showing the full temporal evolution of spatial niche formation rather than initial and final snapshots alone.
